# RHBDL4 protects against endoplasmic reticulum stress by regulating the morphology and distribution of ER sheets

**DOI:** 10.1101/2021.06.15.448480

**Authors:** Viorica L. Lastun, Clémence Levet, Matthew Freeman

## Abstract

In metazoans, the architecture of the endoplasmic reticulum (ER) differs between cell types, and undergoes major changes through the cell cycle and according to physiological needs. Although much is known about how the different ER morphologies are generated and maintained, especially the ER tubules, how context dependent changes in ER shape and distribution are regulated and the factors involved are less characterized. Here, we show that RHBDL4, an ER-resident rhomboid protease, modulates the shape and distribution of the ER, especially under conditions that require rapid changes in the ER sheet distribution, including ER stress. RHBDL4 interacts with CLIMP-63, a protein involved in ER sheet stabilisation, and with the cytoskeleton. Mice lacking RHBDL4 are sensitive to ER stress and develop liver steatosis, a phenotype associated with unresolved ER stress. Our data introduce a new physiological role of RHBDL4 and also imply that this function does not require its enzymatic activity.

## Introduction

The ER is the largest membrane-bound organelle of eukaryotic cells, comprising the nuclear envelope and peripheral ER. In metazoans, the peripheral ER spreads throughout the cytoplasm as a network of interconnected flat sheets and tubules; the flat sheets are present mostly around the nucleus while the tubules interconnect at the level of three-way junctions generating a reticular network towards the cell periphery ^1, 2^. The two distinct ER morphologies accommodate different functions. The sheets are studded with ribosomes and form the rough ER, the site of import, folding and quality control of secreted and transmembrane proteins, and are abundant in professional secretory cells ^3, 4^. The tubules mostly lack ribosomes and form the smooth ER, associated with lipid synthesis and Ca^2+^signalling/homeostasis, abundant in steroid secreting cell ^3–6^. The ER tubules also engage in contacts with other membrane-bound, as well as membraneless, organelles, and a role is emerging for the ER in regulating the biogenesis and/or the dynamics of organelles with which it shares contact sites ^7, 8^.

Several proteins that shape the ER have been discovered. Reticulons and REEPs generate the high membrane curvature characteristic of ER tubules, while Atlastins mediate the homotypic fusion of ER membranes ^9–14^. ER sheets have a more complex structure and less is known about how they are generated and maintained. Reticulons generate the curvature at sheet edges, and several components were shown to stabilise the flat sheets, including the coiled-coil proteins CLIMP-63, p180 and kinectin, the attached polysomes engaged in protein translation, as well as the actin cytoskeleton through myosin 1c ^4, 15, 16^. None of these factors, however, are essential for the existence of ER sheets, suggesting additional structural and regulatory complexity. Beyond its morphology, the factors that contribute to the positioning of the ER within the cell are also not well established; again, more is known about tubules than sheets. ER tubules are often situated along microtubules, and their growth and movement in the cells rely largely on this interaction ^17–19^. ER-sheet proteins CLIMP-63 and p180 can also bind to microtubules, and cytoskeleton depolymerisation was shown to change ER sheet distribution ^15, 20, 21^, although the mechanisms involved in positioning and the dynamics of the ER sheets remain poorly understood. The ER network varies in volume, shape and distribution between cell types and it undergoes reorganisation through the cell cycle, differentiation, and according to physiological needs ^3, 22, 23^, implying the need for regulation. The abundance of ER-shaping proteins is a determining factor of the sheet to tubule ratio: high levels of reticulons and REEP favour the abundance of tubules over the ER sheets, while CLIMP-63 has opposite effect ^4, 24^. Overall, though, rather little is known about the regulation of the rapid and complex changes in ER morphology and distribution in different cellular contexts.

One challenge to ER function, and implicitly to its morphology and distribution, is ER stress. The homeostasis of protein folding in the ER is essential for healthy cells and organisms, and is maintained by a balance between protein biosynthesis and folding on one hand, and export and degradation on the other. Conditions that perturb this homeostasis, generically referred to as ER stress, lead to the accumulation of unfolded proteins in the ER lumen and the activation of the unfolded protein response (UPR). The UPR is an integrated signalling network from the ER to cytoplasm and nucleus, which counteracts the effects of ER stress by upregulating ER chaperones, folding enzymes, and components of the ER-associated protein degradation (ERAD) machinery, while also reducing protein translation ^25, 26^. ER stress also causes significant changes in the size, morphology and distribution of the ER in mammalian and yeast cells ^27, 28^.

RHBDL4 (aka RHBDD1) is a member of the rhomboid intramembrane serine proteases; it is localised in the ER and is upregulated by ER stress ^29, 30^. Although its overall core function remains mysterious, several roles have been assigned to RHBDL4. Its increased expression has been correlated with cancer progression ^31, 32^. RHBDL4 has also been shown to activate EGFR signalling by promoting the secretion in extracellular microvesicles of the growth factor TGFα ^33, 34^. And, in a more classic protease function, RHBDL4 is reported to cleave amyloid precursor protein (APP) ^35^. The most well characterized RHBDL4 function, and the one most apparently relevant to its ER localization, is a function in cleaving and promoting the degradation of various unstable proteins or orphan subunits of protein complexes ^29, 36^, implying that RHBDL4 participates in some forms of ERAD.

Here, we describe an unexpected role of RHBDL4 in regulating the shape and distribution of the ER. By gain and loss of function experiments, we found that the levels of RHBDL4 modulate the shape and distribution of the ER, particularly under ER stress and when the cytoskeleton is disrupted, conditions that require rapid, dynamic changes in ER sheet distribution. We report that RHBDL4 interacts with CLIMP-63 and it potentially associates with the cytoskeleton. Unexpectedly, this role of RHBDL4 appears to be independent of its enzymatic activity, implying an additional, non-proteolytic function of this intramembrane protease. Finally, we also show that RHBDL4 is essential for resolving acute ER stress in mice.

## Results

### 1. The levels of RHBDL4 affect the ER organisation

#### Overexpression of RHBDL4 disrupts ER morphology

RHBDL4 is an ER-resident protein ^29^. While doing transient transfections and immunofluorescence, we noticed that overexpressed RHBDL4 staining looked unusual in many cells. To characterise this within the context of ER organisation, we used CLIMP-63 as a marker for ER sheets and RTN4 for the whole ER – reticulons reside in ER tubules and the edges of the ER sheets ^4^. When expressed at low levels, RHBDL4 showed a perinuclear localisation, characteristic of ER sheets, similar to that of CLIMP-63 (Fig. 1a, second column, the cell indicated by arrowhead). At higher expression levels, RHBDL4 distributed widely into the ER and induced the appearance of punctate structures. Some of the RTN4 was recruited to the puncta, while CLIMP-63 redistributed widely towards the cell periphery into reticular structures (Fig. 1a, second and third column, the cells indicated by arrow). Very high expression levels of RHBDL4 led to a massive disruption in the appearance of the ER (Fig. 1a, fourth column), especially at the level of ER sheets, as shown by CLIMP-63 distribution: part of CLIMP-63 was redistributed into tubules toward the cell periphery, while the rest was confined into compact structures around the nucleus.

**Figure 1.**
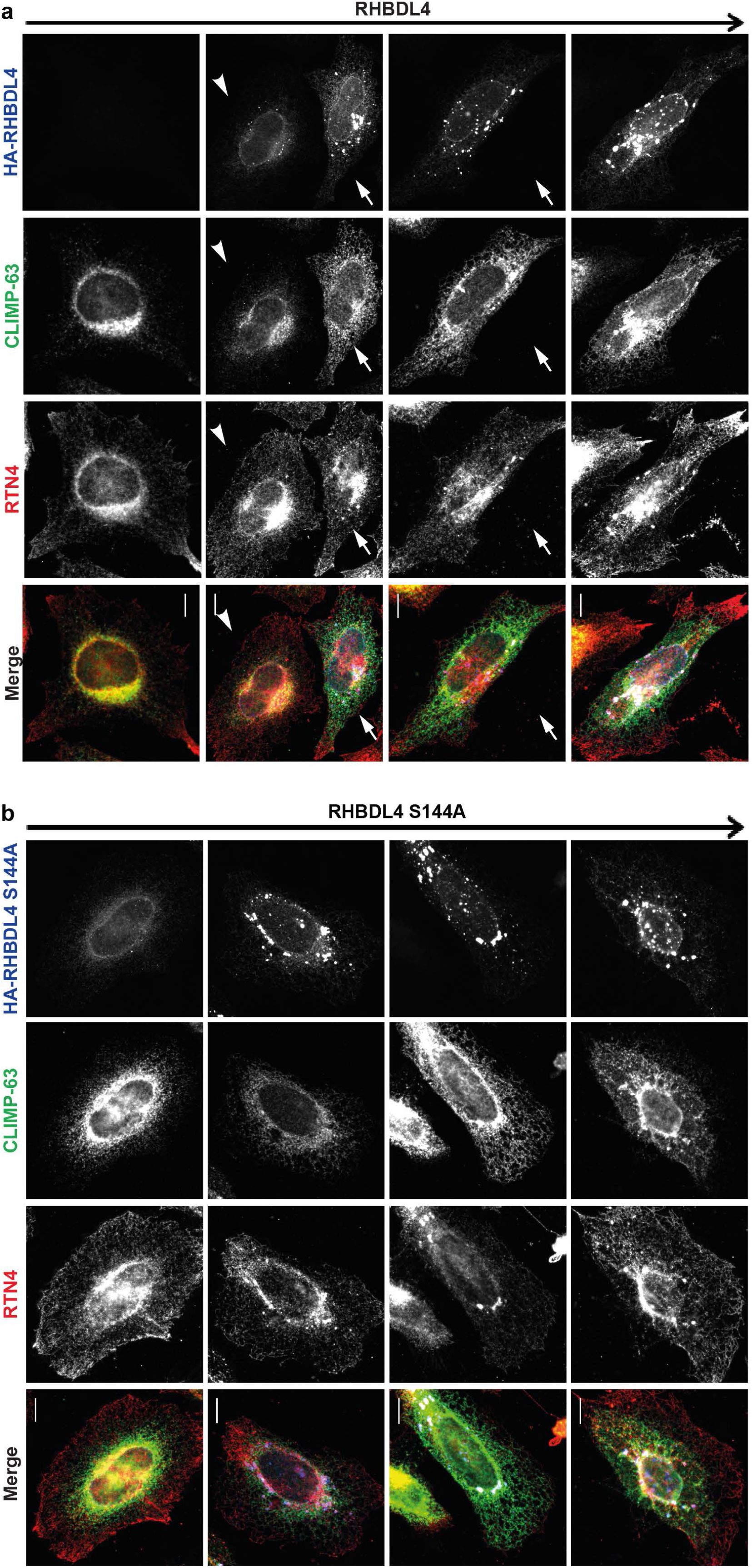
RHBDL4 overexpression disrupts ER organisation. **a**, Immunofluorescence images of RHBDL4 overexpression in HeLa cells: the arrow on the top indicates increasing expression levels. Cells were transfected with HA-tagged RHBDL4 and, after 24 hours, stained for HA tag (blue), RTN4 (red) and CLIMP-63 (green). **b**, Overexpression of HA-tagged RHBDL4 S144A in HeLa cells, analysed as in **a**. Scale bar 10 µm. The images are representative of at least three independent experiments.

Overexpression of a RHBDL4 mutant that had the catalytic serine replaced by an alanine – RHBDL4 S144A ^37^ – showed similar results: increasing levels of RHBDL4 S144A led to increased disruption of ER architecture (Fig. 1b), suggesting that the proteolytic activity of RHBDL4 is not required in this context. This disruption of ER morphology was specific to RHBDL4: the overexpression of another rhomboid protease, RHBDL3, which is structurally very similar to RHBDL4 did not induce changes in the ER morphology, even when expressed at very high levels (Fig. 1S). To make this a more rigorous control for ER-localised RHBDL4, RHBDL3, which normally moves through the ER to its primary plasma membrane location ^30^, was artificially retained in the ER by the addition of a KDEL retention sequence at its C-terminus^38^.

#### RHBDL4 KO affects the ER sheet distribution

Given the effect of RHBDL4 overexpression on the ER, we hypothesized that RHBDL4 plays a role in the ER shape and/or distribution under physiological conditions. To test this, we used two different RHBDL4 KO cell lines – mouse embryonic fibroblasts (MEFs) and human HeLa cells – and compared their ER distribution to that of their wild-type (WT) counterparts.

In WT and RHBDL4 KO MEFs, CLIMP-63 had a similar distribution (Fig. 2a). To confirm this, we quantified the signal distribution using CellProfiler-3.0.0 ^39^. The area between the nucleus and plasma membrane was divided into ten bins (Fig. 2b) and the mean fractional intensity at a given radius was calculated as fraction of total intensity normalized by the fraction of pixels at that radius. Indeed, there was no difference between WT and RHBDL4 KO MEFs (Fig. 2c). The shape of the ER needs to change in response to varying demands on its function. As RHBDL4 was shown to be upregulated in response to ER stress ^29^, we asked whether RHBDL4 is important for the ER shape changes that occur during ER stress. We challenged the cells with tunicamycin, an inhibitor of N-linked glycosylation that leads to accumulation of misfolded proteins in the ER and therefore induces ER stress ^40, 41^. In WT MEFs, after 24 hours of tunicamycin treatment, CLIMP-63 redistributed dramatically throughout the cytoplasm as compared to the control (Fig. 2d, upper row), implying a major reorganisation of ER sheets. In contrast, CLIMP-63 redistribution was abnormal in RHDBL4 KO MEFs, with 28.76% of cells showing a strong phenotype, with a “collapse” of CLIMP-63 signal around the nucleus (Fig. 2d, lower row, and quantified in Fig. 2e). This phenotype was significantly rescued by transient transfection of RHBDL4 WT (Fig. 2e), indicating that the effect was specific to loss of RHBDL4. We could not detect rescue when using the RHBDL4 S144A mutant (data not shown) but this result is hard to interpret because expression of this mutant, even at low levels, induces acute ER stress on its own ^29^ (Fig. S2, lanes 3 and 4 compared to lanes 7 and 8). It is therefore likely that its expression combined with tunicamycin treatment induced very high levels of ER stress in RHBDL4 KO cells, preventing a rescue of ER morphology.

**Figure 2.**
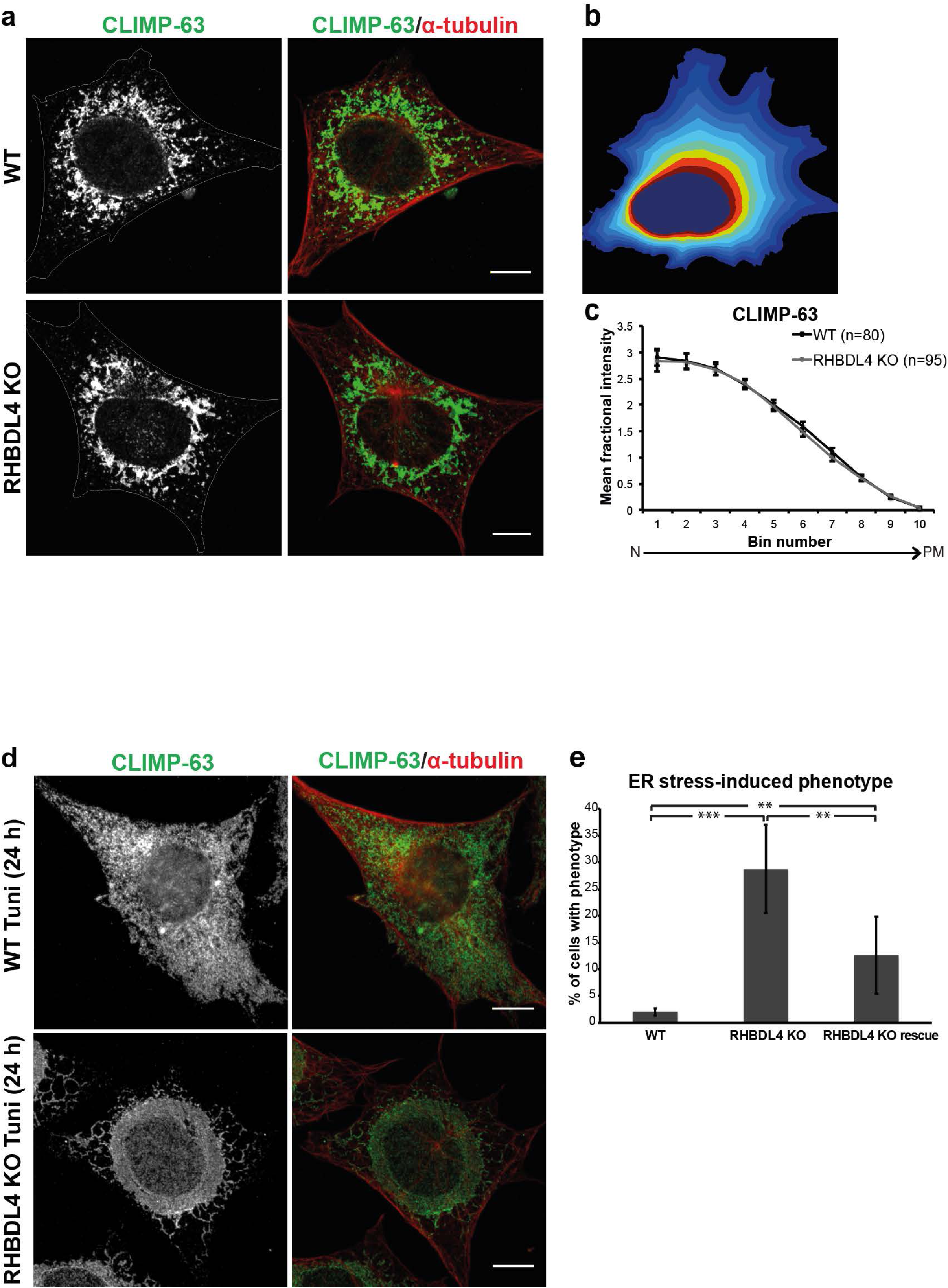
RHBDL4 affects ER distribution under ER stress conditions in MEFs. **a**, Immunofluorescence of CLIMP-63 (green) and α-tubulin (red) in WT and RHBDL4 KO MEFs. Scale bar 10 µm. **b**, A representation of the way the area between the nucleus and plasma membrane was divided into ten bins to quantify signal distribution. **c**, Quantification of CLIMP-63 distribution. Error bars represent the 95% CI. **d**, As in **a,** but 24 hours after tunicamycin treatment. The images and quantification are representative of three independent experiments. **e**, Quantification of cells presenting the tunicamycin-induced phenotype. For rescue experiments, RHBDL4 KO cells were transiently transfected with HA-tagged RHBDL4 WT. Error bars represent 95% CI (n=3 experiments, with at least 50 cells/experiment analysed for WT and RHBDL4 KO and at least 25 cells/experiment analysed for rescue RHBDL4-HA). For WT vs. RHBDL4 KO, p=0.0002 t-test, two sided (three stars on the graph) and p=0.049 Mann-Whitney-Wilcoxon test, two sided. For rescue RHBDL4-HA vs. RHBDL4 KO, p=0.0033 t-test (two stars on the graph) and p=0.049 Mann-Whitney-Wilcoxon test, two sided. For rescue RHBDL4-HA vs. WT, p=0.0033 t-test (two stars on the graph) and p=0.049 Mann-Whitney-Wilcoxon test, two sided.

In WT HeLa cells, the distribution of CLIMP-63 and RTN4 followed their expected localisation: CLIMP-63 was more concentrated within the innermost bins, while RTN4 was distributed more widely towards the plasma membrane (Fig 3a, upper row, and quantified in Fig. S3). The ER was more compacted around the nucleus in the RHBDL4 KO cells when compared to the WT, especially when looking at CLIMP-63 (Fig. 3a, lower row, and quantified in Fig. 3b and 3c). Transient transfection of the RHBDL4 KO cells with either WT or the catalytically inactive RHBDL4 rescued the phenotype (Fig. 3d), supporting the conclusion that the role of RHBDL4 in ER sheet distribution is not dependent on its proteolytic activity.

**Figure 3.**
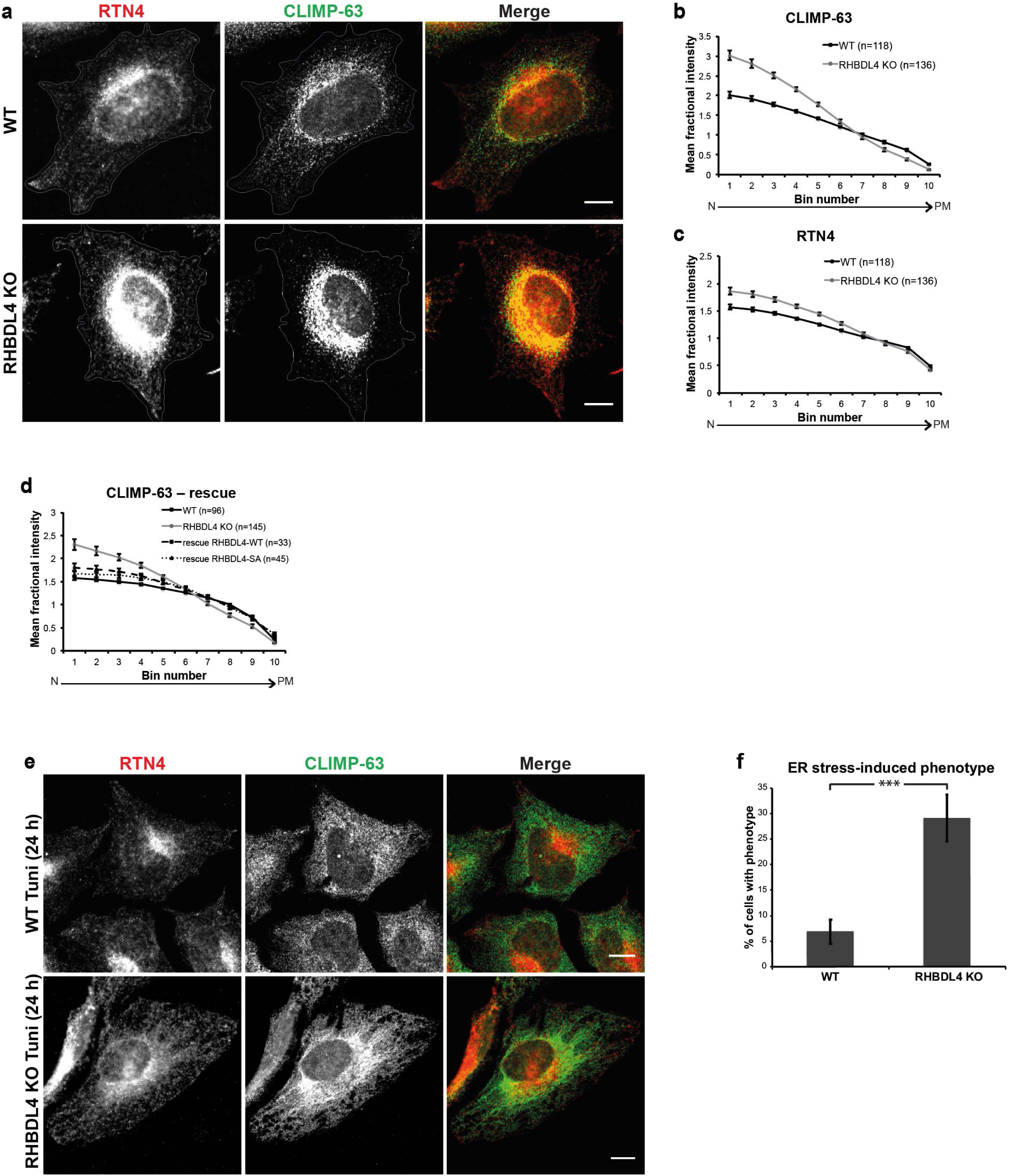
RHBDL4 affects ER distribution in HeLa cells. **a**, Immunofluorescence of RTN4 (red) and CLIMP-63 (green) in WT and RHBDL4 KO HeLa cells. **b** and **c,** Comparison of CLIMP-63 (**b**) and RTN4 (**c**) distribution between WT and RHBDL4 KO HeLa cells. Error bars represent the 95% CI. **d**, Rescue of the phenotype. Quantification of CLIMP-63 distribution in RHBDL4 KO cells transiently transfected with HA-tagged RHBDL4 WT or RHBDL4 S144A. Error bars represent the 95% CI. **e,** Immunofluorescence of CLIMP-63 and RTN4 in WT and RHBDL4 KO cells 24 hours after the tunicamycin treatment. **f**, Quantification of cells presenting the tunicamycin-induced phenotype. Error bars represent the 95% CI (n=4 experiments, with over 100 cells/experiment analysed). p<0.0001 t-test, two sided (three stars on the graph) and p=0.021 Mann-Whitney-Wilcoxon test, two sided. The images and quantifications are representative of at least three independent experiments.

As in MEFs, tunicamycin treatment of HeLa cells led to a redistribution of CLIMP-63 throughout the cytoplasm in WT cells, whereas, in RHDBL4 KO, a subset of cells showed a pronounced phenotype of more concentrated perinuclear staining and a non-homogeneous distribution towards the cell periphery (Fig. 3e, 3f).

Altogether our results in MEFs and HeLa cells show that RHBDL4 is important for ER sheet distribution and that the effect was more pronounced during ER stress.

#### The levels of ER shaping proteins and the ER stress response are similar in WT and RHBDL4 KO cells

The role of RHBDL4 in ER shape/distribution could be direct or indirect, via other ER-shaping proteins. Also, the phenotypes we described in RHBDL4 KO cells could be due to a defective folding/secretory function of the ER, or failure to activate the UPR and cope with the ER stress. Analysis of MEFs and HeLa cells by western blot showed no obvious differences in the levels of RTN4, ATL1 or CLIMP-63 between WT and RHBDL4 KO, neither in control nor under induced ER stress conditions (Fig. 4a and Fig. S4a, respectively). This implies that RHBDL4 does not affect the levels of these ER-shaping proteins.

**Figure 4.**
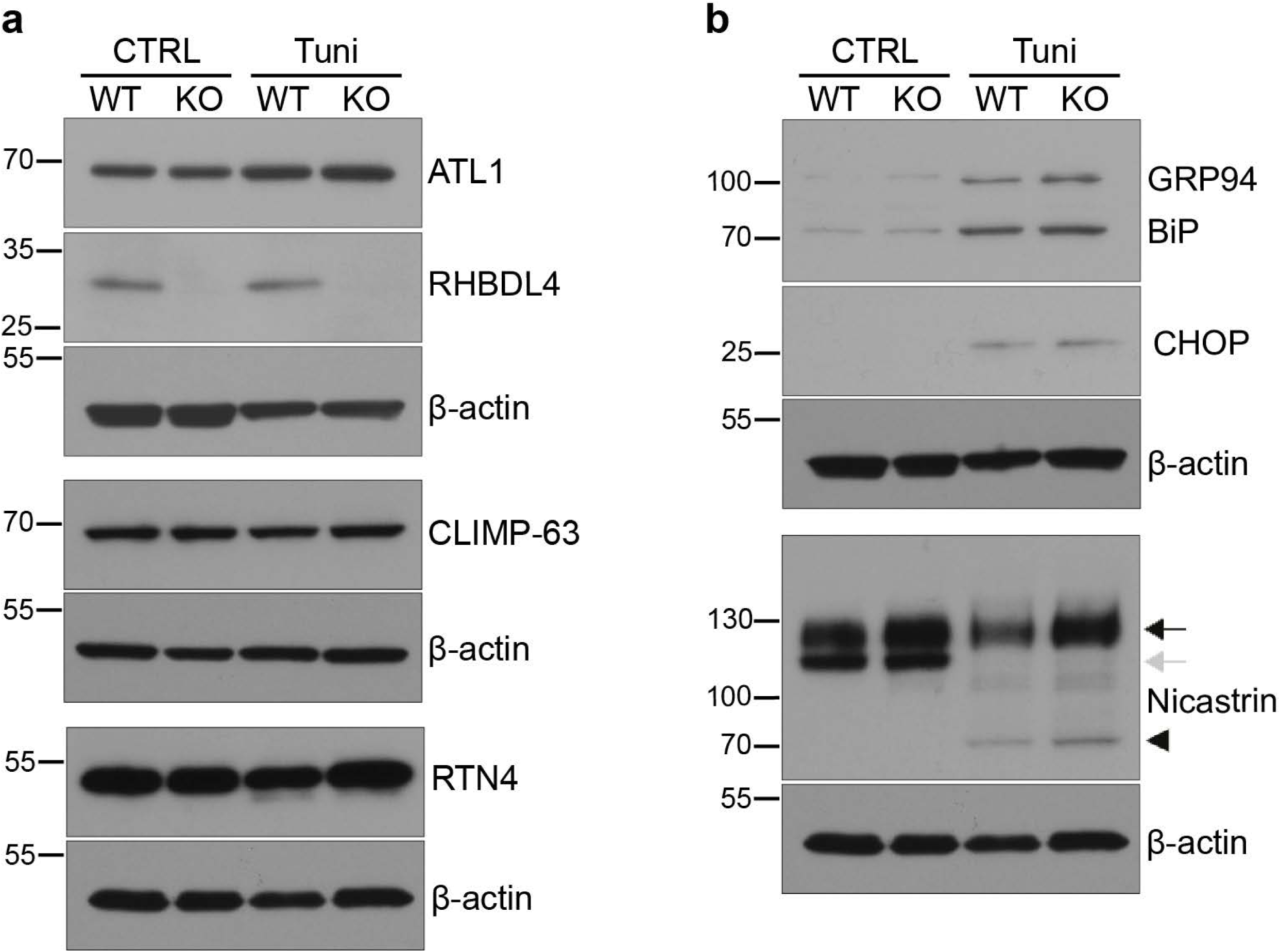
The levels of ER shaping proteins and the ER stress response are similar in WT and RHBDL4 KO cells. Western blot of total cell lysates from WT and RHBDL4 KO MEFs in control conditions or 24 hours after tunicamycin treatment. **a,** Western blots of RHBDL4 and ER shaping proteins ATL1, CLIMP-63 and RTN. **b,** Western blots of UPR targets BiP and GRP94 (detected using an anti-KDEL antibody), CHOP, and Nicastrin. The black arrow represents the mature post-Golgi protein, while the grey arrow represents the immature, ER-localised one. The arrowhead represents the non-glycosylated Nicastrin, visible in the tunicamycin-treated samples. β-Actin was used as a loading control and it was developed on the stripped membranes, previously used for the proteins above each β-actin panel. The results are representative of three independent experiments.

We next asked whether RHBDL4 affects the ER stress response. Under ER stress, to cope with the folding load, cells increase their protein folding capacity by activating the UPR ^42^. Two of the chaperones that are upregulated most in response to ER stress are BiP and GRP94 ^43^. As shown in Figure 4b and Fig S4b, BiP and GRP94 upregulation by Tunicamycin treatment was similar in WT and RHBDL4 KO cells. Failure to restore protein homeostasis in the ER results in activation of apoptosis. We assayed a central mediator of apoptosis, the pro-apoptotic transcription factor CHOP ^44^, and found similar levels in WT and RHBDL4 KO MEFs (Fig. 4b). We conclude that the ER stress response is not significantly affected in these RHBDL4 KO cells.

To assess the general secretory function of the ER in KO cells, we examined the cellular location of Nicastrin, a component of the γ-secretase complex, which is found in two forms: one immature, ER-localised, detected as a lower molecular weight band in western blots, and one mature, post-Golgi, detected as a higher molecular weight form ^45^. We found no differences in the levels of post-Golgi Nicastrin, neither in control nor under ER stress conditions (Fig. 4b and Fig S4b). The efficiency of tunicamycin treatment was similar in the WT and RHBDL4 KO cells, as shown by the appearance of non-glycosylated species of Nicastrin. This implies that the overall secretory function of RHBDL4 KO cells is not significantly compromised. Taking all these results together, we conclude that the role of RHBDL4 in shaping the ER under stress conditions is not a secondary effect of a role in homeostasis of ER shaping proteins, response to ER stress, or general secretion.

### 2. RHBDL4 localises to the ER sheets and interacts with CLIMP-63

The phenotypes we observed so far indicate that RHBDL4 levels have an impact on the ER sheets, so we wanted to determine the localization of endogenous RHBDL4 within the ER. Several commercial antibodies as well as one produced by our group failed to detect the endogenous RHBDL4 by immunofluorescence. We therefore used a biochemical approach to examine endogenous RHBDL4 localisation. We isolated microsomes from mouse liver as well as from HeLa cells, and used differential centrifugation to separate the rough and smooth ER on a sucrose cushion ^46, 47^. As sketched in Figure 5a, left, following ultracentrifugation the rough ER (ER sheets studded with ribosomes) sediments through the 1.3 M sucrose cushion, while the smooth ER (ER tubules) segregates at the interface between the two sucrose concentrations; the material in between is a rough/smooth, or intermediate, ER ^5^. For HeLa cells, due to a lower amount of material, the smooth ER was not readily visible and, therefore, we analysed all fractions starting one millilitre above the interface and finishing with the pellet, as indicated in Figure 5a, right side.

**Figure 5.**
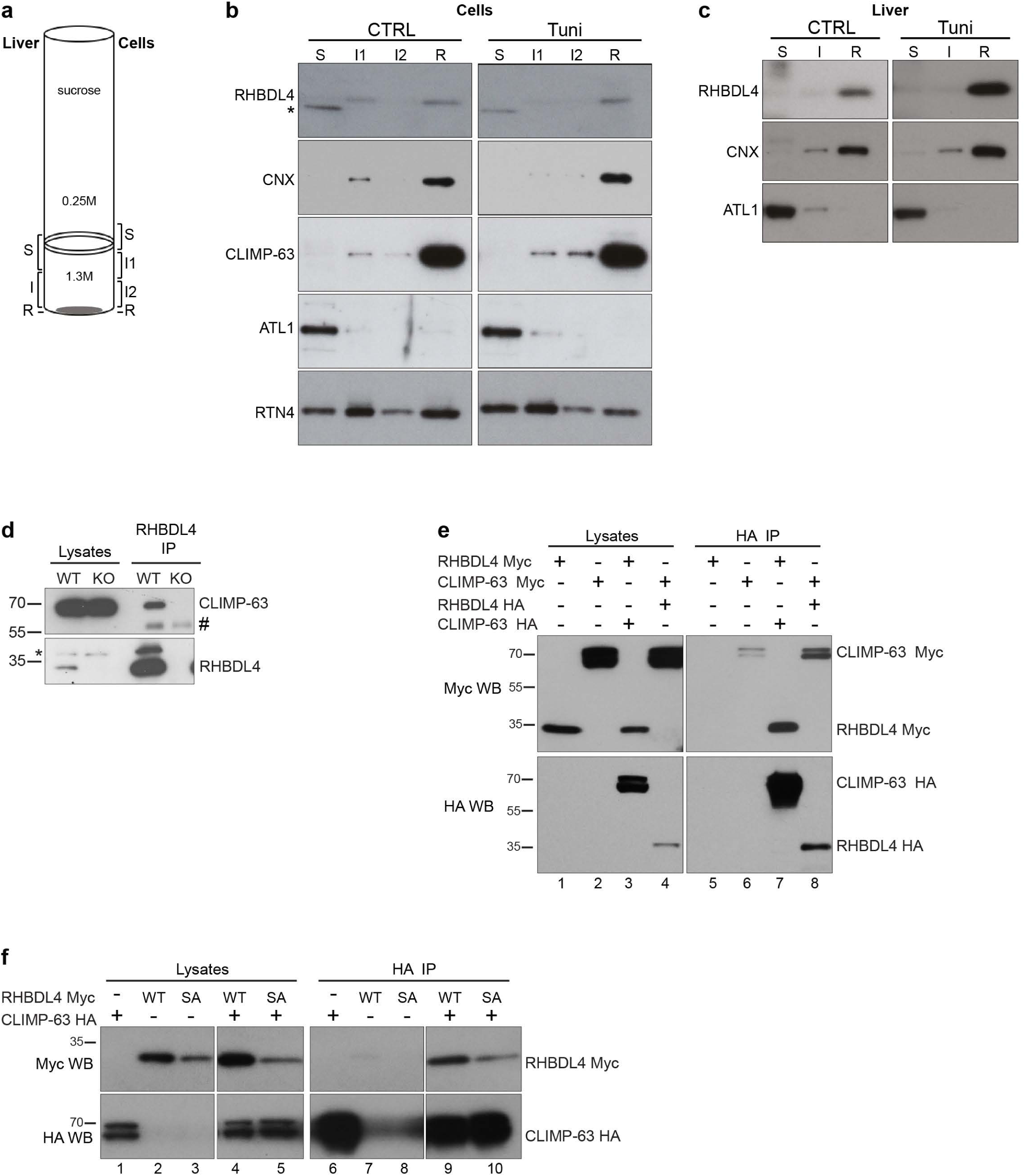
RHBDL4 resides within the ER sheets and interacts with CLIMP-63. **a**, Separation of rough and smooth ER by differential centrifugation – schematic representation. **b,** Microsomes from HeLa cells, control and tunicamycin treated, were separated as in **a**. Smooth ER (S), rough ER (R) and intermediate ER were analysed by western blot for the indicated ER proteins. The asterisk represents a non-specific band. The results are representative of three independent experiments. **c**, Microsomes from mouse liver, control and tunicamycin treated, were separated as indicated in **a** and analysed as in **b**. The results are representative of three mice per condition. **d**, Immunoprecipitation of endogenous RHBDL4 followed by western blot for CLIMP-63 and RHBDL4. The # represents IgG heavy chain and asterisk a non-specific band. **e**, Immunoprecipitation of tagged versions of RHBDL4 WT and CLIMP-63, transiently expressed in HeLa cells, followed by western blot as indicated. **f**, As in **e,** but including both the WT and S144A (SA) mutant RHBDL4. The results are representative of three independent experiments.

In agreement with their published localisation, by this fractionation method we found ATL1 located in the smooth ER, RTN4 in both smooth and rough ER, while Calnexin (CNX) – an ER-resident chaperone – and CLIMP-63 were specific to the rough ER (Fig. 5b and 5c), ^4, 48^, confirming a good separation. In both HeLa cells and mouse liver, RHBDL4 distributed predominantly to the rough ER (Fig. 5b and 5c). Upon ER stress there was no obvious change in RHBDL4 distribution in mouse liver, and a somewhat increased level in rough ER in HeLa cells. We obtained similar results in MEFs (Fig. S5).

These results show that endogenous RHBDL4 is localised predominantly within the ER sheets, which is consistent with the ER disruption phenotypes we observed.

Next, we asked whether we could detect interactions between RHBDL4 and ER shaping proteins. In co-immunoprecipitation experiments, we found that endogenous RHBDL4 physically interacts with CLIMP-63 (Fig. 5d). Significantly, this interaction was also found in our previous large-scale interaction screen of the RHBDL4 interactome – CLIMP-63 aka CKAP4 was one of the reproducible hits in HeLa and HEK 293 cells ^49^. We could not detect an interaction between RHBDL4 and RTN4 or ATL1, or between RHBDL4 and CNX, another ER sheet membrane protein (not shown). The interaction between endogenous RHBDL4 and CLIMP-63 was recapitulated using overexpressed, tagged proteins: CLIMP-63 HA pulled down RHBDL4 Myc and, in turn, RHBDL4 HA pulled down CLIMP-63 Myc (Fig. 5e, lane 7 and 8, respectively). Both, RHBDL4 WT and RHBDL4 S144A co-immunoprecipitated with CLIMP-63 (Fig. 5f, lanes 9 and 10), indicating that the interaction with CLIMP-63 did not depend on the RHBDL4 active site.

### 3. RHBDL4 associates with the cytoskeleton

CLIMP-63 interacts with microtubules ^20^, and given the impact of RHBDL4 overexpression and KO on the shape and distribution of the ER, we asked whether RHBDL4 might also interact with the cytoskeleton, which provides a functional scaffold for shaping and distributing the ER.

Using an approach expected to preserve protein-cytoskeleton interactions, we separated the cytoskeletal fraction from HeLa cells ^50^. During the fractionation, the actin cytoskeleton is preserved, as well as the cold-stable microtubules often described as the ones that interact with other proteins ^51^ (Fig. 6, the lower two rows show actin and tubulin). Strikingly, most of RHBDL4 and CLIMP-63 segregated into the cytoskeletal fraction while RTN4 and ATL1 did not (Fig. 6a, lane 3). CLIMP-63 was present into the cytoskeletal fraction independent of RHBDL4 (Fig. 6a, lane 6 compared to lane 3). Likewise, RHBDL4 segregated into the cytoskeletal fraction regardless the amounts of CLIMP-63 (Fig. 6b, lane 6 compared to lane 3), suggesting that the two proteins associate with the cytoskeleton independently of each other. The cytoskeletal association of HA-RHBDL4 did not depend on its proteolytic activity, as shown by the analysis of transiently expressed HA-RHBDL4 WT and HA-RHBDL4 S144A (Fig. 6c, lanes 6 and 9, respectively); to exclude any interference of endogenous RHBDL4, these experiments were done in RHBDL4 KO cells. Similar results were obtained in MEFs (Fig. S6).

**Figure 6.**
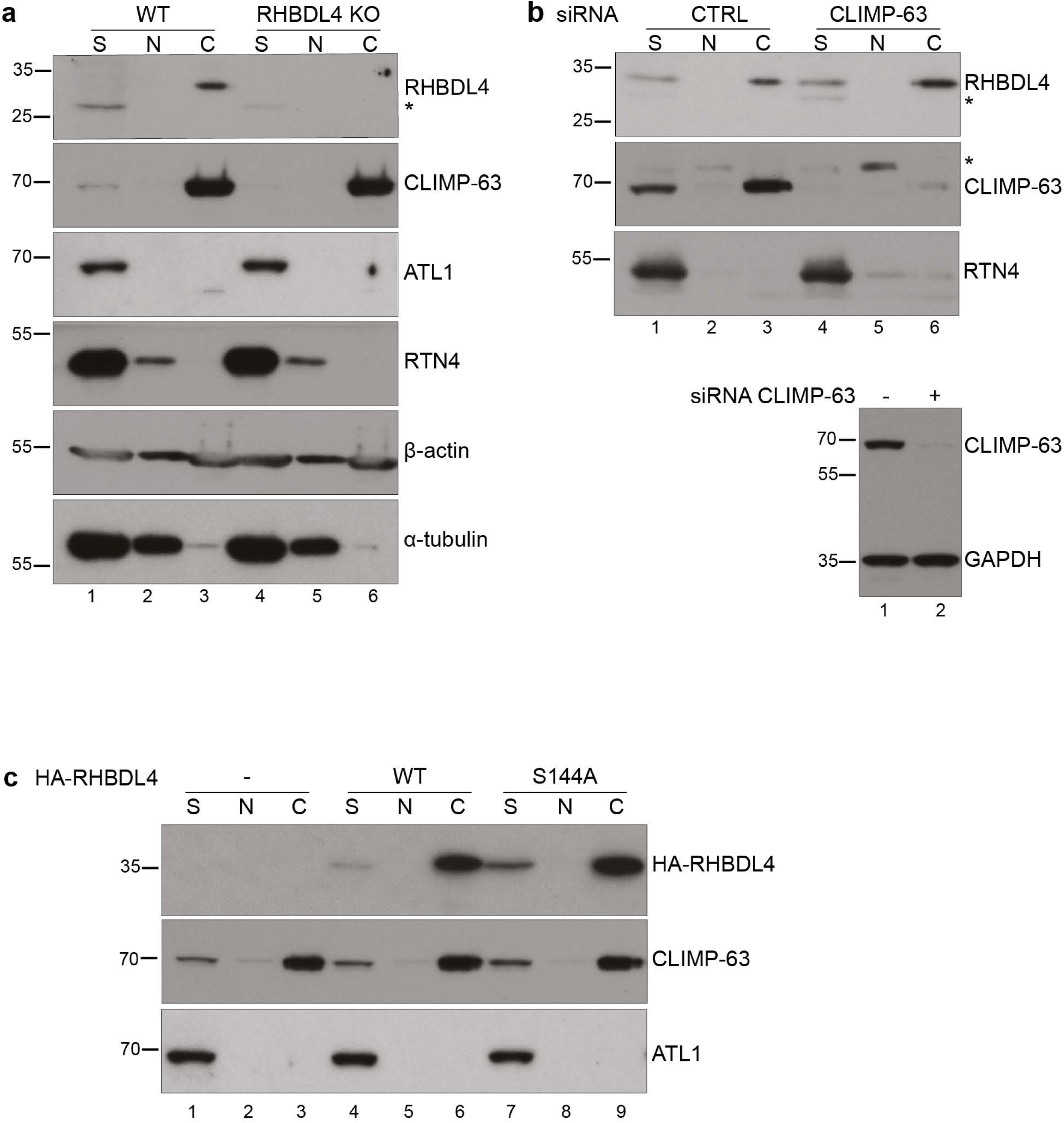
RHBDL4 associates with the cytoskeleton. **a,** Western blot of soluble (S), nuclear (N) and cytoskeletal (C) fractions isolated from WT and RHBDL4 KO HeLa cells and analysed for ER proteins – RHBDL4, CNX, CLIMP-63, ATL1, RTN4 – as well as for β-actin and ⍺-tubulin. **b**, Top panel as in **a**, western blot of S, N and C fractions isolated from HeLa cells treated with CLIMP-63 siRNA 72 hours before analysis. In the lower panel are shown the levels of CLIMP-63 in total cell lysates from the same experiment. **c**, Anti-HA tag western blot of C, N and S fractions isolated from RHBDL4 KO HeLa cells transiently expressing HA-tagged RHBDL4 WT or RHBDL4 S144A. In **a** and **b**, the asterisk represents non-specific bands. The results are representative of three independent experiments.

When overexpressed, RHBDL4 disrupted the microtubule organization in a concentration-dependent manner: cells expressing low/moderate levels of RHBDL4 had normal microtubule morphology, while cells with higher expression showed dramatically reorganized microtubules (Fig. 7a, second versus third and fourth column). Overexpressing the RHBDL4 S144A mutant showed similar results (Fig. 7b).

**Figure 7.**
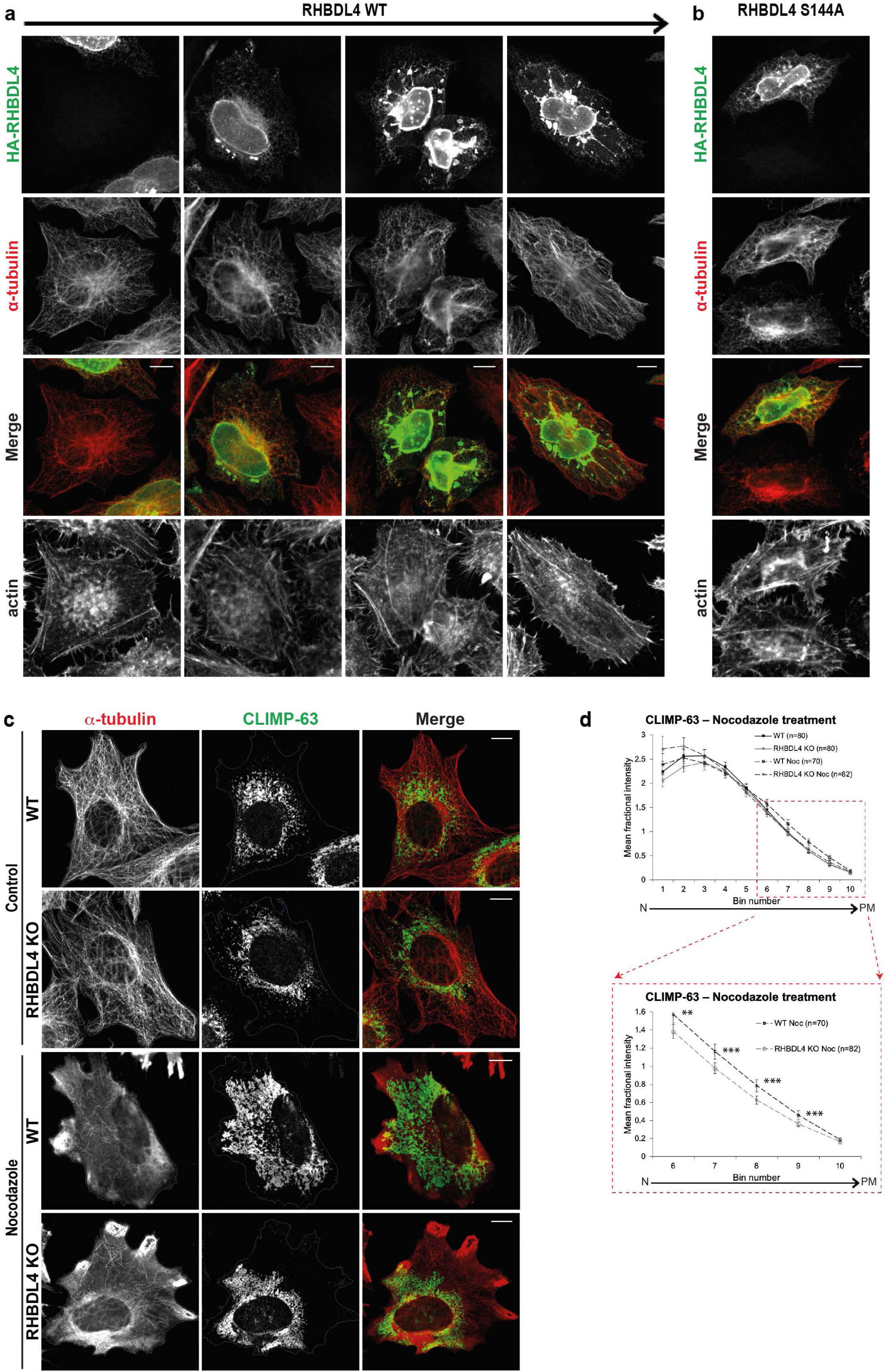
RHBDL4 and microtubules. Immunoflourescence of HeLa cells transiently transfected with HA-tagged RHBDL4 WT in **a,** or RHBDL4 S144A in **b**. Cells were stained for HA tag (green), α-tubulin (red) and actin (lower row). The arrow at the top indicates increasing expression levels. The images are representative of three independent experiments. **c**, Immunoflourescence of WT and RHBDL4 KO MEFs showing the microtubule (α-tubulin – red) and ER-sheet (CLIMP-63 – green) distribution under control or nocodazole treatment. **d**, Quantification of CLIMP-63 distribution (left) and an enlarged area of the graph to emphasise the statistical significance in the peripheral bins (right). Error bars represent the 95% CI. p=0.0027 t-test (two stars on the graph) and p=0.0092 Mann-Whitney-Wilcoxon test, two sided, for bin 6; p=0.0004 t-test (three stars on the graph) and 0.0029 Mann-Whitney-Wilcoxon test, two sided, for bin 7; p=0.0001 t-test (three stars on the graph) and 0.0003 Mann-Whitney-Wilcoxon test, two sided, for bin 8; p=0.0007 t-test (three stars on the graph) and 0.0015 Mann-Whitney-Wilcoxon test, two sided for bin 9; p=0.0297 t-test (one stars on the graph) and p=0.0211 Mann-Whitney-Wilcoxon test, two sided for bin 10. The number of cells analysed is shown on the graph. For **c** and **d** the results are representative of two independent experiments.

Microtubule depolymerization results in rapid reorganisation of the ER network ^17, 52, 53^. We used this to investigate further a role for RHBDL4 in ER dynamics and distribution in the context of ER-cytoskeleton interaction. We exposed WT and RHBDL4 KO MEFs and HeLa cells to nocodazole, to disrupt the microtubules, and analysed the resulting changes in ER sheet morphology and distribution. In WT cells, both MEFs and HeLa, the nocodazole treatment led to wide spreading of CLIMP-63 signal throughout the cytoplasm, confirming that the ER sheets underwent a major reorganisation (Fig. 7c, third row and Fig. S7a). In RHBDL4 KO, CLIMP-63 redistribution was less pronounced and, in many cells there were peripheral areas devoid of ER sheets (Fig. 7c, fourth row and Fig S7a). Quantification of this CLIMP-63 spreading phenotype confirmed the difference in the peripheral distribution between WT and RHBDL4 KO cells (Fig. 7d, f and Fig. S7b). These results showed that, as when ER stress was induced by tunicamycin treatment, in the absence of RHBDL4, microtubule-dependent ER sheet dynamics are defective.

### 4. RHBDL4 protects against ER stress in vivo

To study the role of RHBDL4 *in vivo* we created RHBDL4 KO mice. We detected no obvious phenotype under normal conditions but the fact that RHBDL4 KO MEFs only showed a phenotype when challenged with tunicamycin (or nocodazole) prompted us to examine the effect of ER stress in KO mice.

We used intraperitoneal injection of a sublethal dose of tunicamycin (1 μg/g body weight) ^54, 55^ to induce ER stress, and followed the mice over three days. The first difference we noticed between WT and RHBDL4 KO mice was in weight loss: at day three of treatment the RHBDL4 KO mice had lost significantly more weight than WT animals (Fig. 8a), suggesting that the former were more severely affected by induced ER stress. Liver is the mouse tissue with the strongest response to tunicamycin-induced ER stress ^56^ and one of the tissues with highest RHBDL4 expression (Fig. 8b). Therefore, it was the tissue of choice to look for further differences between the WT and RHBDL4 KO mice. Upon dissection, we also noticed a striking difference in the appearance of WT and RHBDL4 KO livers at three days of treatment, with the latter being pale in colour, a sign of liver steatosis. Oil Red O staining of liver sections confirmed this; after three days of treatment, the RHBDL4 KO mice accumulated dramatically elevated levels of neutral lipids in their livers when compared to WT (Fig. 8c). At day one, there was no apparent difference between the two genotypes (Fig. S8).

**Figure 8.**
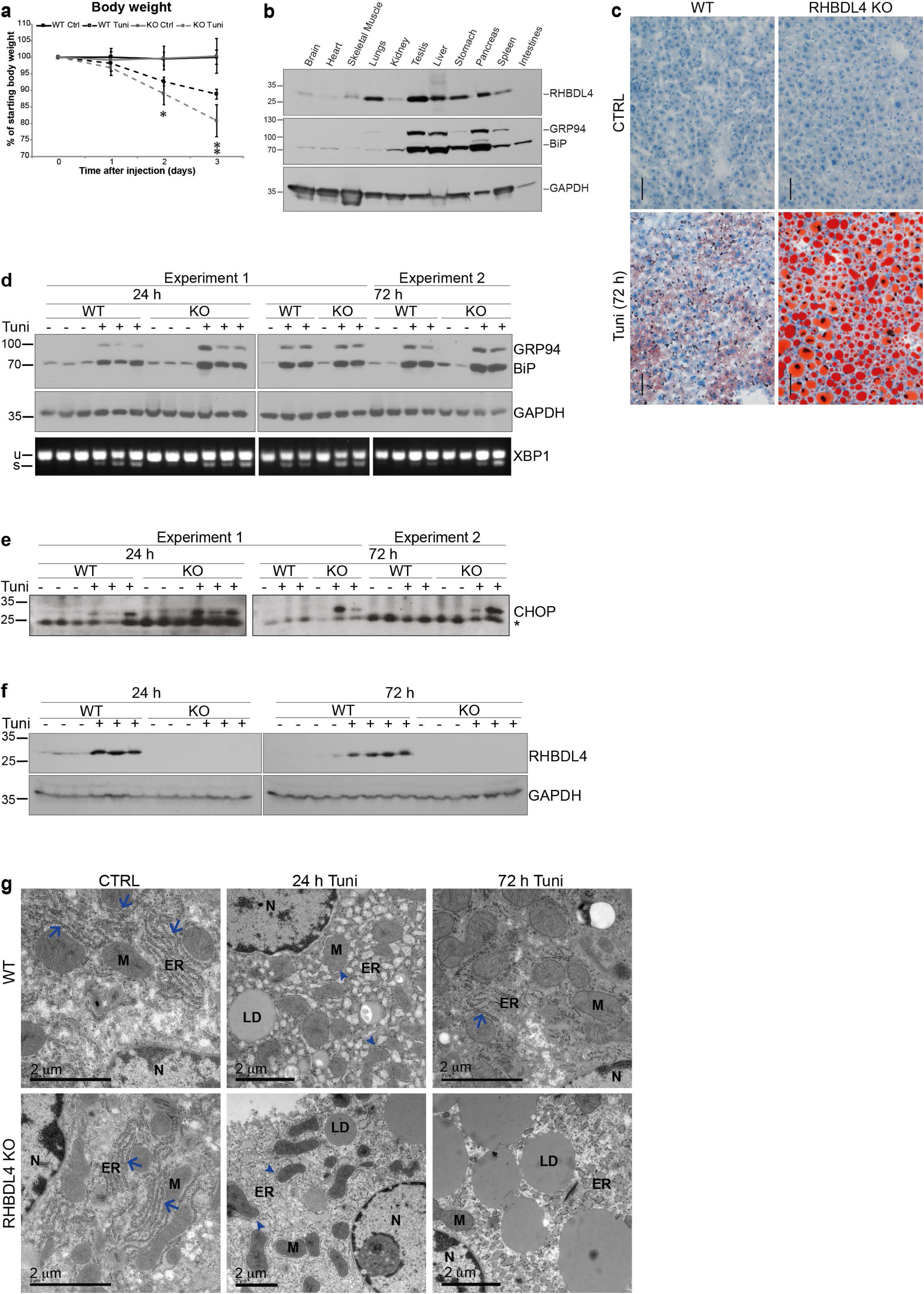
RHBDL4 protects against the ER stress in mice. **a,** Body weight changes of WT and RHBDL4 KO mice control or treated with tunicamycin. Errors bars represent 95% CI (six to eight mice per genotype per condition, combined from the three different experiments). The asterisks represent the statistical significance for WT Tuni vs. RHBDL4 KO Tuni: day 1, p=0.3324, t-test, two sided, and p=0.4875 Mann-Whitney-Wilcoxon test; day 2, p=0.0230, t-test, two sided (one star on the graph), and p=0.0151 Mann-Whitney-Wilcoxon test; day 3 p=0.0014 t-test, two sided (two stars on the graph), and p=0.0151 Mann-Whitney-Wilcoxon test, two sided. **b,** Western blot for RHBDL4 in mouse liver lysates from WT mice. BiP and GRP94 (detected using an anti-KDEL antibody) were used as ER markers, and GAPDH was used as a loading control. **c,** Oil red O staining of liver tissue sections from WT and RHBDL4 KO mice, control or 72 hours after tunicamycin treatment. The results are representative of two experiments with a total of four to six mice per genotype per condition. **d,** Western blot for BiP and GRP94 (detected using an anti-KDEL antibody) in mouse liver lysates from WT and RHBDL4 KO mice, control or tunicamycin treated for the indicated times (top panels). GAPDH was used as loading control. The lower panels show the corresponding XBP1 splicing, examined by RT-PCR using primers flanking the intron region. **e**, Western blot for CHOP. The asterisk represents a non-specific band which serves as a loading control. **f**, Western blot as in **d,** but for RHBDL4. **g,** Electron microscopy of liver tissue sections from WT and RHBDL4 KO mice, control, 24 or 72 hours after tunicamycin treatment. N, nucleus; M, mitochondria; ER, endoplasmic reticulum; LD, lipid droplet. The results are representative of at least three mice per genotype per condition.

To check whether the failure of RHBDL4 KO mice to cope with ER stress induced by tunicamycin was due to a deficient activation of the UPR, we measured the upregulation of UPR targets BiP and GRP94, as well as another marker of UPR activation, splicing of the X-box binding protein-1 (XBP1). In response to ER stress, XBP1 mRNA undergoes splicing and is translated into an active transcription factor. The transcriptional program triggered by XBP1 leads to ER expansion, upregulation of ERAD and secretory pathway components ^57^. We found no differences between WT and RHBDL4 KO mice (Fig. 8d), indicating that RHBDL4 KO mice do not have a defect in UPR activation. Nevertheless, three days after tunicamycin treatment, the ER stress was unresolved in RHBDL4 KO mice, as indicated by the persistence of the pro-apoptotic transcription factor CHOP (Fig. 8e). RHBDL4 expression itself was upregulated by the tunicamycin treatment, consistent with a role for it in coping to ER stress (Fig. 8f).

For a better insight into the phenotype of RHBDL4 KO mice under ER stress, we performed transmission electron microscopy (TEM) on liver sections and looked at the appearance of the ER. Under control conditions, the ER looked similar in WT and RHBDL4 KO livers, with stacks of rough ER packed around the mitochondria (Fig. 8g, left column, blue arrows). Twenty-four hours after tunicamycin treatment, in both genotypes the rough ER lost the appearance of organised stacks, became dilated and distributed broadly throughout the cytoplasm, with only scarce ER sheets around mitochondria (Fig. 8g, middle column). By 72 hours after tunicamycin treatment, a clear difference between the genotypes became clear: in WT livers the ER had started to recover into structures of stacks around the mitochondria, while in the RHBDL4 KO the rough ER remained dilated, filling the cytoplasm, and large lipid droplets were accumulated (Fig. 8g, right column). These TEM data further demonstrate a major defect in the resolution of ER stress in the absence of RHBDL4.

## Discussion

In metazoans, the ER has a complex architecture with different ER morphologies hosting different functions. The ER architecture differs between cell types and undergoes changes during the cell cycle and according to physiological needs. There is significant knowledge about how the ER tubules are generated and maintained, but not as much is known about the ER sheets. How do rapid changes in shape and distribution of the organelle take place and what are the regulators; what factors contribute to the positioning of the ER within the cell, and how does the ER interact with the cytoskeleton? We report here that RHBDL4 plays a role in ER sheet dynamics during ER stress, that it interacts with CLIMP-63 and the cytoskeleton, and that it is essential for coping with ER stress in mice.

We have shown that endogenous RHBDL4 resides primarily within the ER sheets. Consistent with this, the phenotypes induced by its overexpression or knockout were mostly at the level of ER sheets, especially during conditions that require their reorganisation/redistribution – ER stress or microtubule depolymerisation. In response to ER stress, mammalian and yeast ER undergoes significant changes in size, morphology and distribution ^27, 28, 57^. In both cell lines we used in this study, human HeLa cells and mouse embryonic fibroblasts, RHBDL4 knockout led to defective redistribution of ER sheets. At the same time, the function of the ER, in terms of secretion, and the activation of the UPR, were similar in WT and RHBDL4 KO cells, suggesting that RHBDL4 participates in adapting ER shape to the cellular context rather than to the ER function itself. Overall, the common theme that emerges from our results is that RHBDL4 has a role in ER sheet dynamics; in its absence, the ER sheets were more compact around the nucleus and more stable to disassembly and redistribution, while moderate overexpression had the opposite effect.

While the reticular network of ER tubules can be reconstituted *in vitro* using a small number of proteins – reticulons, REEPs and Atlastins ^10, 11^ – the ER sheets have a more complex structure, and it is not clear how they are generated and maintained. None of the components shown to stabilize ER sheets is essential, and even though there are indications that the ER sheets interact with both microtubules and the actin cytoskeleton ^15, 20^, the actual mechanisms involved in their dynamics and positioning are poorly understood. We found an interaction between RHBDL4 and the ER sheet protein CLIMP-63. Our fractionation experiments demonstrated that both proteins segregated into the cytoskeleton binding fraction, although it is important to note that their interaction with the cytoskeleton was independent of each other, suggesting that RHBDL4 associates with the cytoskeleton either directly or via a different intermediate. When overexpressed, CLIMP-63 co-aligns with microtubules and together they from bundles around the nucleus, thus disrupting the microtubule organization ^20^ (data not shown). Overexpression of RHBDL4 disrupted microtubule organization too, but the effect was different to that produced by CLIMP-63, consistent with their independent interaction with microtubules. CLIMP-63 forms homo-oligomers via its luminal domain, which limit its movement within the plane of membrane ^58^. Also, CLIMP-63 is thought to form intermembrane bridges to stabilize the ER sheets ^4, 59^. One possibility is that RHBDL4 interferes with the formation of CLIMP-63 homo-oligomers or intraluminal bridges to allow ER sheets to be dynamic when needed. We emphasise that this is a speculative idea at this stage but overall our data suggest that, while CLIMP-63 plays a structural role in ER sheets, RHBDL4 appears to be involved in their dynamics, playing a regulatory role.

To date, there is no reported substrate for RHBDL4 that could explain this role and none of the ER shaping proteins we checked appeared to be cleaved by RHBDL4. Indeed, the effects we describe here using overexpression were similar for RHBDL4 WT and the catalytically inactive mutant RHBDL4 S144A: the effect on ER and microtubules, the segregation into the cytoskeletal fraction and the interaction with CLIMP-63. Moreover, both RHBDL4 WT and RHBDL4 S144A rescued partially the phenotype in RHBDL4 KO HeLa in control conditions. This leads us to conclude that the function we describe in ER shape and dynamics is independent of the catalytic function of this rhomboid protease. This conclusion is made more complex by the fact that expression of RHBDL4 S144A induces ER stress, making the interpretation of some experiments difficult. Even though we used low expression levels for rescue experiments, the ER stress induced by expressing the mutant can mask a rescue of RHBDL4 function. Nevertheless, the balance of all the evidence in both cell types suggests a nonproteolytic function for RHBDL4 in this context. To our knowledge, this is the first report of a nonproteolytic function for a functional secretase rhomboid (i.e. not an iRhom or other more distant member of the rhomboid-like superfamily ^60^), however it is not new in the world of enzymes. Among enzymes shown to fulfil additional non-catalytic functions are kinases, matrix metalloproteinases (MMPs) and presenilins, and the non-catalytic functions of these enzymes range from scaffolding to protein trafficking and calcium signalling ^61–64^.

Pharmacological ER stress induces transient lipid droplet (LD) accumulation in mouse liver and, when combined with a defective UPR signalling, it leads to hepatic steatosis. Genetic disruption of any of the three UPR branches or of protein quality control results in failure to cope with the ER stress, and hepatic steatosis ^28, 55^, most likely caused by a failure to upregulate the necessary folding machinery. When challenged with tunicamycin, RHBDL4 KO mice showed a phenotype similar to that described for UPR deficient mice: significant weight loss and liver steatosis, characterised by massive accumulation of LDs. However, in RHBDL4 KO mice, UPR activation was normal (as shown by XBP1 splicing, and BiP and GRP94 upregulation), suggesting a different cause for the failure to cope with the ER stress. Three days after tunicamycin treatment, in WT livers, ER sheets began to reassemble around the mitochondria, whereas in RHBDL4 KO livers, the ER remained dilated and scattered through the cytoplasm, and large LDs were common. Although ER stress results in LD accumulation, and the disruption of LD biogenesis induces the UPR ^65^, the role of LDs in ER stress response remains unclear. Our data further emphasise the relationship between ER stress and LD accumulation but further work is needed to uncover the factors and mechanisms involved in ER sheet reassembly and LD clearing after ER stress: our work suggests that RHBDL4 participates in this stress resolution. Notably, some proteins with well defined roles in ER shape – for example, Atlastin, REEP1 and Spastin – are known to affect LD biology ^66–68^, but none of these roles have been studied in the context of ER stress.

In conclusion, while the relevant molecular and cellular mechanisms remain to be discovered, we have uncovered a physiological role for RHBDL4 in regulating the morphology and distribution of ER sheets. Our work has also shown that this previously unrecognised RHBDL4 function is essential in cells for coping with conditions that require dynamic reorganisation of ER sheets, and in mice for resolving the pathology associated with ER stress.

## Methods

### Cell culture, DNA constructs and transfections

MEFs and HeLa cells (WT and RHBDL4KO) were cultured in Dulbecco’s Modified Eagle’s Medium (DMEM) supplemented with 10% heat inactivated Fetal Bovine Serum (FBS) and penicillin-streptomycin, at 37°C in 5% CO_2_. The triple HA-tagged RHBDL4 construct in pcDNA3.1 plasmid has been described previously ^33^. Using an untagged version of this plasmid, a C-terminal Myc tag was inserted using site-directed mutagenesis. CLIMP-63 cDNA was purchased from Open Biosystems (MHS6278-202833153), subcloned into pcDNA3.1, and HA or Myc tag were added by site-directed mutagenesis. All constructs were validated by sequencing. For overexpression and rescue experiments, 60-70% confluent cells were transfected with the cDNA of interest in pcDNA3.1 plasmid. Polyethylenimine (PEI, 25 kDa linear, Polysciences) mediated transfections were carried out using a DNA to PEI ratio of 1:2.5 (w/w), in complete medium, for HeLa cells. For MEFs, a DNA to PEI ratio of 1:3 (w/w) was used, and the transfections were done in serum free medium for 5 hours, after which the transfection medium was replaced with complete, fresh DMEM. The cells were analysed 24 hours post transfection.

### Generation of MEFs

Embryonic fibroblasts were generated from RHBDL4 WT and RHBDL4 KO E14.5 embryos and immortalized using lentiviral transduction of SV40 virus large T antigen (Ef1a_Large T-antigen_Ires_Puro, Addgene plasmid 18922), as described by Christova et al. (2013) ^69^.

### RHBDL4 KO HeLa cells

The RHBDL4 KO HeLa cells were generated using the CRISPR/Cas9 system. The gRNAs were designed online (http://www.e-crisp.org) ^70^. Four different gRNAs were cloned into pSpCas9(BB)-2A-Puro (PX459) plasmid ^71^ and tested in HeLa cells. The cells were transfected with PX459 plasmid containing the gRNAs and, 48 hours post transfection, were treated with puromycin to select the transfectants. The levels of RHBDL4 were assessed by Western blot and the most efficient gRNA (TAGTGTGGAGAAGTGTTACC, targeting the first coding exon) was selected for further use in this study. Single cell clones transfected with the above gRNA were selected from and used.

### U2OS Flp-In stable cell lines

U2OS Flp-In stable cell lines expressing N-terminal triple HA-tagged RHBDL4 (WT or S144A) were obtained by co-transfecting the pcDNA5/FRT vector containing the cDNA for RHBDL4, and the pOG44 plasmid at a ratio of 1:9. Stably transfected cells were selected and maintained in DMEM supplemented with 10% FBS and 200 µg/ml Hygromycin. To induce the expression of RHBDL4, ∼70% confluent cells were treated with tetracycline and analysed by Western blot 24 hours later.

### Tunicamycin and Nocodazole treatment

To induce ER stress, the cell culture medium of ∼70% confluent cells was replaced with fresh medium containing tunicamycin (0.1 µg/ml for MEFs and 5 µg/ml for HeLa, final concentration, T7765; Sigma-Aldrich), and the cells were incubated at 37°C, 5% CO_2_, for 24 hours. For microtubule depolymerisation experiments, the cell culture medium of ∼70% confluent cells was replaced with fresh medium containing 10 μM Nocodazole (M1404; Sigma-Aldrich) and cells were incubated at 37°C, 5% CO_2_, for 15 minutes. In both cases, control cells were treated with DMSO.

### Immunoprecipitation

All experiments were performed on ∼80% confluent cells. For endogenous RHBDL4, the immunoprecipitation was done using the Pierce™ Crosslink Magnetic IP/Co-IP Kit (88805; Thermo Scientific), according to the manufacturer’s protocol. The cells were washed twice with ice-cold phosphate-buffered saline (PBS) and lysed in lysis buffer (25mM Tris, 150mM NaCl, 1mM EDTA, 1% NP40, 5% glycerol, pH 7.4) containing EDTA-free protease inhibitor cocktail (11873580001; Roche). Lysates were centrifuged at 10,000g for 10 min and the post-nuclear supernatants were used for subsequent immunoprecipitation. Cell lysates were incubated with a rabbit anti-RHBDL4 antibody, described by Fleig et al. (2012)^29^, crosslinked to Protein A/G magnetic beads for 1 hour at 4°C on a rotator. Immunoprecipitates were washed three times with lysis buffer and eluted with the included elution buffer (pH 2.0). The eluates were neutralised, mixed with 5X sample buffer including 50 mM DTT and incubated at 65°C for 10 min prior loading onto gels. Cell lysates were prepared with 5X sample buffer and DTT, and incubated at 65°C, alongside the eluates. Samples were analysed by SDS-PAGE (Novex, WedgeWell 8-16% Tris-Glycine Mini Gels) followed by Western blot. For overexpressed RHBDL4 and CLIMP-63, immunoprecipitation was done similarly to the above, except the lysis buffer contained CHAPS instead of NP40, and anti-HA or anti-Myc magnetic beads were used.

### Cytoskeleton Enrichment

The cytoskeletal fraction was separated using the ProteoExtract® Cytoskeleton Enrichment and Isolation Kit (17-10195; Millipore), following the manufacturer’s protocol. The experiments were performed on 80-90% confluent cells. At the end of collection, the soluble, nuclear and cytoskeletal fractions were adjusted to equal volumes and mixed with 5x SDS-PAGE sample buffer and 50 nM DTT. The samples were incubated at 65°C for 10 min and equal volumes were loaded onto gels (Novex, WedgeWell 8-16% Tris-Glycine Mini Gels) and analysed by Western blot.

### Microsome separation and ER fractionation

Microsomes were separated from MEFs, HeLa cells or mouse liver using the protocol described by Song et al. (2006) ^47^, adapted. All steps of the protocol were performed on ice and all solutions contained protease inhibitor cocktails. The livers from sacrificed animals were rapidly removed and placed in ice-cold PBS. After three washes with PBS, the livers were cut into pieces and weighed. Small pieces (1-3 mm^3^) were mixed with five volumes (w/v) of homogenisation solution (0.25 M sucrose; 10 mM HEPES; pH 7.5) and homogenised by five manual strokes using a Dounce tissue grinder, loose-fitting pestle. The homogenates were centrifuged at 1,000g, 4°C, for 10 minutes. The supernatants were collected and centrifuged at 8,000g, 4°C, for 15 minutes. The supernatants were diluted with homogenisation solution and 15 mM CsCl to a total volume of 6 ml, and laid gently on a cushion of 1.3 M sucrose, 10 mM HEPES, 15 mM CsCl, pH 7.5, and centrifuged at 200,000g for 2 hours, at 4°C. One ml fractions starting from the interface (0.25 M/1.3 M sucrose) were collected. The pellet was solubilised in 1 ml of 0.25 M sucrose, 10 mM HEPES, 15 mM CsCl, pH 7.5. The fractions were mixed with 5X SDS-PAGE sample buffer and 50 mM DTT and incubated at 65°C for 10 min. Equal volumes were loaded onto gels (Novex, WedgeWell 8-16% Tris-Glycine Mini Gels) and analysed by Western blot.

Microsome separation from cells was done using the same protocol, except the homogenisation was done using the tight-fitting pestle and 25 strokes, and the fractions were collected starting 1 ml above the interface. For each sample, three 10-cm dishes of 80-90% confluent cells were used. The cells were washed one time with ice-cold PBS and one time with homogenisation buffer before being scraped in homogenisation buffer and transferred to the Dounce tissue grinder.

### Western blot

For Figure 4 and Figure S4, the cells were lysed in ice-cold RIPA buffer (150 mM sodium chloride, Triton X-100, 0.5% sodium deoxycholate, 0.1% SDS, 50 mM Tris, pH 8.0), containing EDTA-free protease inhibitor cocktail and Benzonase (E1014-5KU; Sigma-Aldrich), after which the lysates were mixed with 5X sample buffer, 50 mM DTT. The whole cell lysates were incubated at 65°C for 10 min prior loading onto gels (Novex, WedgeWell 8-16% Tris-Glycine Mini Gels). For Western blot analysis of mouse tissues, small pieces of tissues were homogenised in RIPA buffer supplemented with protease inhibitor cocktail, using a handheld tissue homogeniser. The homogenates were centrifuged at 10,000g for 15 minutes and supernatants were mixed with 5X sample buffer and 50 mM DTT and incubated at 65°C for 10 min prior loading onto gels (Novex, WedgeWell 8-16% Tris-Glycine Mini Gels). For all other experiments, the sample preparation is described in the corresponding Methods sections. After electrophoresis, the proteins were transferred to polyvinylidene difluoride (PVDF) membrane (Millipore). Membranes were blocked in 5 % low fat milk in PBS and then incubated with primary antibodies in 0.1 % Tween-PBS at 4°C overnight, for anti-CHOP, or at room temperature for one hour, for the rest of the antibodies. After three washes, membranes were incubated with the appropriate HRP-conjugated secondary antibodies for 30 min and washed three times. Antibodies were visualized using Amersham ECL detection reagent (RPN2209) and X-ray films. The following primary and secondary antibodies were used: rabbit polyclonal anti-RHBDL4 obtained as described by Fleig et al. (2012)^29^, rat monoclonal anti-HA-HRP (Roche, 12 013 819 001), mouse monoclonal anti-beta-Actin-HRP (Sigma-Aldrich, A3854), rabbit anti-ATL1 (Santa Cruz sc-67232), mouse anti-RTN4 (Nogo, Santa Cruz, sc-271878), mouse anti-KDEL (Abcam, ab12223), mouse anti-alpha-Tubulin (Abcam, ab-7291), rabbit anti-CLIMP-63 (CKAP4, Proteintech, 16686-1-AP), goat anti-Nicastrin (Santa Cruz sc-14369), mouse anti-CHOP (Cell Signalling 2895), rabbit anti-Calnexin (Santa Cruz sc-11397), goat anti-Myc-HRP (Abcam, ab1261), mouse anti-GAPDH (Sigma, G8795), mouse anti-vimentin (kit # 17-10195, Millipore), goat polyclonal anti-rabbit-HRP (Biorad, 170-6515), mouse monoclonal anti-goat-HRP (Santa Cruz, sc-2354) or goat polyclonal anti-mouse-HRP (Santa Cruz, sc-2055).

### Immunofluorescence

For immunofluorescence, MEFs and HeLa cells were grown in 8-well culture slides uncoated or coated with Poly-D-Lysine, respectively (Falcon, 354108, and Corning BioCoat, 354632). Before staining, cells were washed twice with PBS, fixed with 4% paraformaldehyde (Electron Microscopy Sciences) in PBS for for 20 min, permeabilised with 0.2% Triton X-100 in PBS for 5 min, and blocked with 1% BSA in PBS for 30 minutes. Cells were incubated with primary antibodies for one hour and with secondary antibodies for 30 minutes, both in blocking buffer. Every step of the above protocol was followed by three PBS washes. Primary antibodies used for staining were: rat anti-HA (Roche 11 867 423 001); rabbit anti-CLIMP-63 (CKAP4, Proteintech, 16686-1-AP), goat anti-RTN4 (Nogo, Santa Cruz sc-11027), mouse anti-alpha-Tubulin (Abcam ab-7291). The fluorophore-conjugated secondary antibodies Alexa Fluor 488, Alexa Fluor 568 and Alexa Fluor 647 donkey anti-mouse, donkey anti-rabbit and donkey anti-goat (all from Invitrogen) were used in various combinations. For actin detection, Alexa Fluor 568 or Alexa Fluor 647 Phalloidin was used. After the last PBS wash, the chamber slides were disassembled, and coverslips were mounted on top of the cells using Fluoromount-G mounting solution containing DAPI (Southern Biotech). Single slice images were taken on a Zeiss 880 microscope using the confocal settings.

### Image analysis and quantification

For figures, the brightness/contrast of images were adjusted in Fiji, using the same settings across images belonging to the same experiment. Quantification of signal intensity was done using CellProfiller -3.0.0 ^39^. The area between the nucleus and plasma membrane was divided into ten bins and the mean fractional intensity at a given radius was calculated as fraction of total intensity normalized by the fraction of pixels at that radius. For cell segmentation, the nuclei were used as primary objects and merged images of Phalloidin or tubulin (for whole cell staining) and CLIMP-63 (ER staining) were used to identify the whole cell area. The cells that were not segmented correctly were manually removed from the analysis.

### Reverse transcriptase polymerase chain reaction (RT-PCR)

To detect the XBP1 splicing, the RNA from mouse liver was isolated using RNeasy Mini Kit (Qiagen, 74104) according to the manufacturer’s instructions. RT-PCR was performed using 0.5 µg total RNA per reaction, 0.6 µM each XBP1 specific primers and one-step RT-PCR (QIAGEN OneStep RT-PCR, 210212). The PCR products were analysed by 2% agarose gel. The primers used were: 5’gaagagaaccacaaactcca 3’ (FW) and 5’ggatatcagactcagaatct 3’.

### Mice

RHBDL4 KO mice were obtained using the Cre-loxP system. Briefly, the targeting construct contained the floxed exon 2 (the first coding exon, encoding the first 144 amino acids, including the catalytic serine) of RHBDL4. The loxP sites were introduced by recombineering and the successful targeting of EK.CCE embryonic stem (ES) cells was confirmed by southern blotting ^72^. Exon 2 was excised in the targeted ES cells by expressing Cre recombinase. Two independent clones of targeted ES cells with excised exon 2 were injected into blastocysts derived from C57BL6 mice and implanted into pseudo-pregnant female ICR mice using standard techniques. The resulting chimeric mice were crossed to WT mice to confirm the germline transmission and to further obtain homozygous RHBDL4 -/- mice in the next generation. Genotyping was performed by PCR and the mice were age and sex matched for experiments.

For tunicamycin-induced ER stress, mice between 9 and 14 weeks old were used across experiments. Tunicamycin stock solution was dissolved in DMSO and diluted 100X in 150 mM sucrose prior injection. One intraperitoneal injection of tunicamycin (1 μg/g body weight) or vehicle was administered, and the mice were followed over the indicated times. Mice were housed in temperature-controlled conditions with a 12-hour light-dark cycle, and fed on standard rodent chow. All procedures were conducted under a UK project licence (PPL 80/2584).

### Mouse liver histology

For Transmission Electron Microscopy (TEM), small pieces of liver (1-2 mm^3^) from sacrificed animals were rapidly collected and transferred to fixative solution (4% PFA + 2.5% glutaraldehyde in 0.1M PIPES pH 7.2) and stored at 4 °C overnight before processing. A Leica EM TP automated processing unit was used for sample processing. Samples were washed three times in 0.1M PIPES buffer, one time in 0.1 M PIPES containing 50 nM glycine to block free aldehydes, followed by a final wash in 0.1M PIPES. Samples were stained with 1% osmium tetroxide in 0.1M PIPES for 2 hours at 4 °C and subsequently washed with four times with water and stained with 0.5% uranyl acetate at 4 °C for approximately 12 hours. After two washes with water, the samples were taken through a dehydration series of 30%, 50%, 70%, 90%, 95% ethanol, 15min each, at room temperature, and three incubations in 100% ethanol, 30 min each. The samples were infiltrated with TAAB low viscosity epoxy resin as it follows: 25% resin in ethanol for 1 hour, 50% resin in ethanol for 3 hours, 75% resin in ethanol for 2 hours, then nine times in 100% resin, 5 hours each. Samples were removed from the tissue processor and placed in Beem capsule molds containing fresh resin, and polymerised at 60 °C for 48 hours. Ultrathin (90 nm) resin sections were obtained using a Diatome diamond knife with a Leica UC7 ultramicrotome and transferred to 200 mesh copper grids. The grids were post-stained for 5 min with Reynold’s lead citrate and imaged using a Gatan Oneview camera in an FEI Tecnai 12 TEM operated at 120kV.

For Oil Red O staining, fresh liver pieces were collected in OCT compound in cryomolds and frozen on dry ice. Cryosections, 10 µm thick, were stained with Oil Red O, and imaged on a COOLSCOPE slide scanner.

### Statistical analysis

The error bars in all figures represent the 95% confidence interval. The statistical significance was calculated using two-sided t-test and two-sided Mann-Whitney-Wilcoxon test.

## Acknowledgements

We thank Sonia Muliyil for critical reading of the manuscript. We thank Yonka Christova for generating the knockout mice and MEFs; Pedro Carvalho for the gift of U2-OS Tet-On cells; Errin Johnson and Anna Pielach (Dunn School EM Facility) for the EM; and Richard Stillion for the ORO staining. Confocal microscopy imaging was done at the Dunn School Bioimaging Facility (Micron Advanced Bioimaging Unit). This work was supported by Wellcome Trust Investigator Awards to MF (101035/Z/13/Z and 220887/Z/20/Z).

## Supplementary figure legends

**Figure S1.**
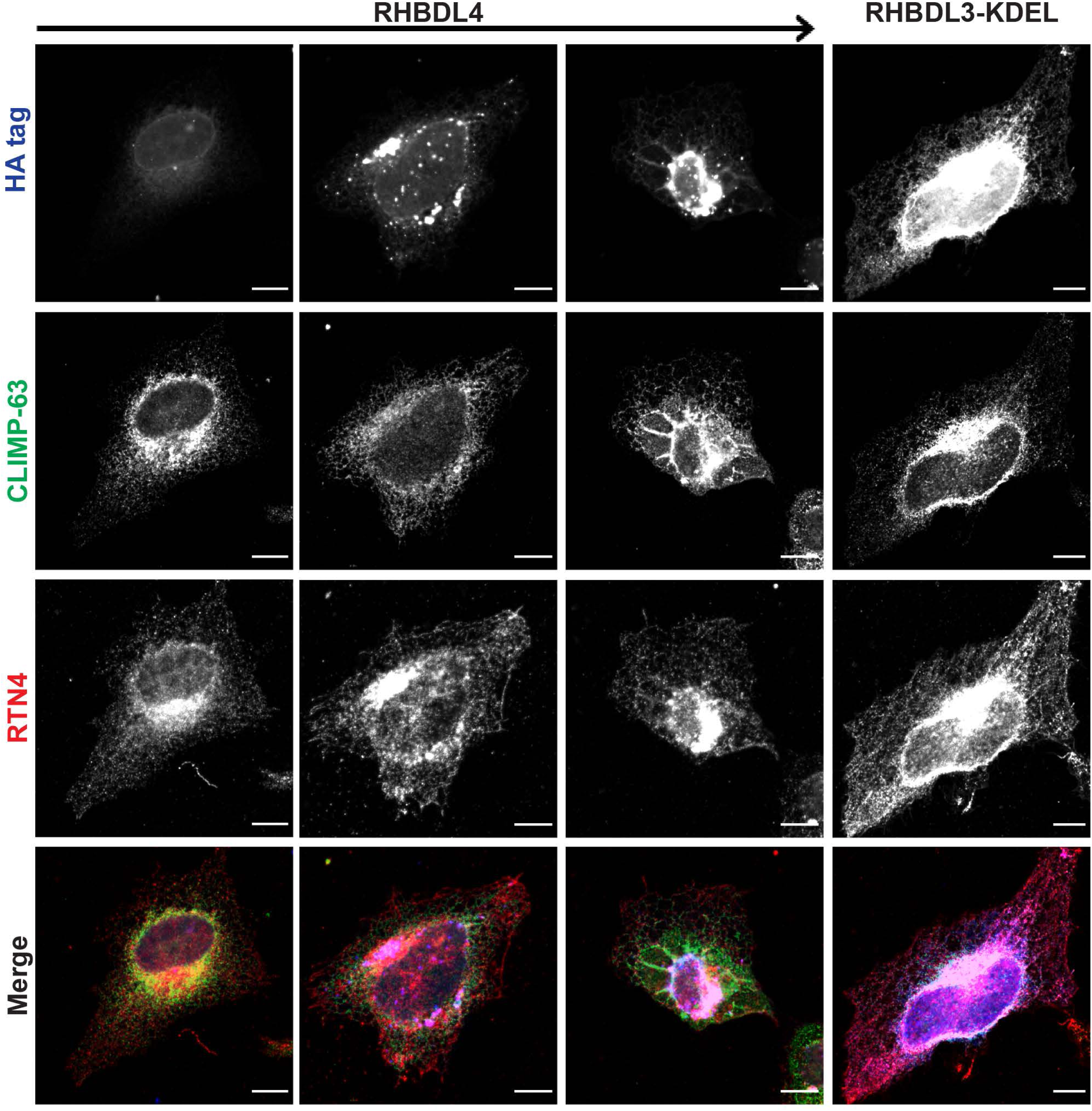
RHBDL4 overexpression disrupts the ER organisation. **a**, Immunofluorescence images of RHBDL4 (left) and RHBDL3-KDEL (right) overexpression in HeLa cells: the arrow on the top indicates increasing expression levels. Cells were transfected with HA-tagged RHBDL4 or RHBDL3-KDEL and, after 24 hours, stained for HA tag (blue), RTN4 (red) and CLIMP-63 (green). Scale bar 10 µm. The images are representative of three independent experiments.

**Figure S2.**
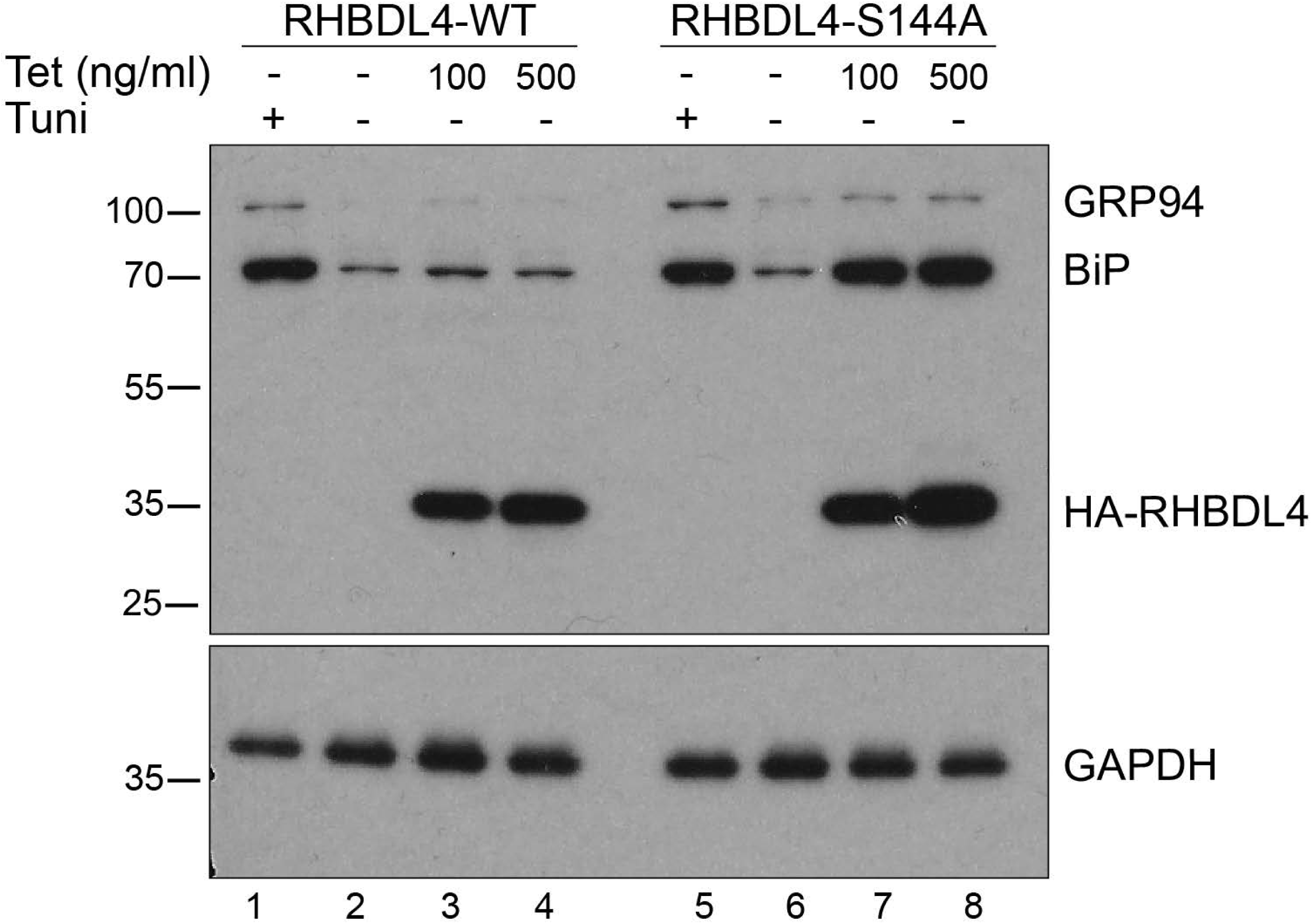
The expression of catalytically inactive RHBDL4 induces ER stress and triggers the UPR. Western blots of UPR targets BiP and GRP94 (detected using an anti-KDEL antibody) in total cell lysates of U2-OS Tet-On cells expressing RHBDL4 WT or RHBDL4 S144A. As a positive control for UPR induction, cells were treated with tunicamycin for 24 hours. The levels of RHBDL4 WT and RHBDL4 S144A were detected using anti-HA antibodies. GAPDH was used as a loading control. The results are representative of three independent experiments.

**Figure S3.**
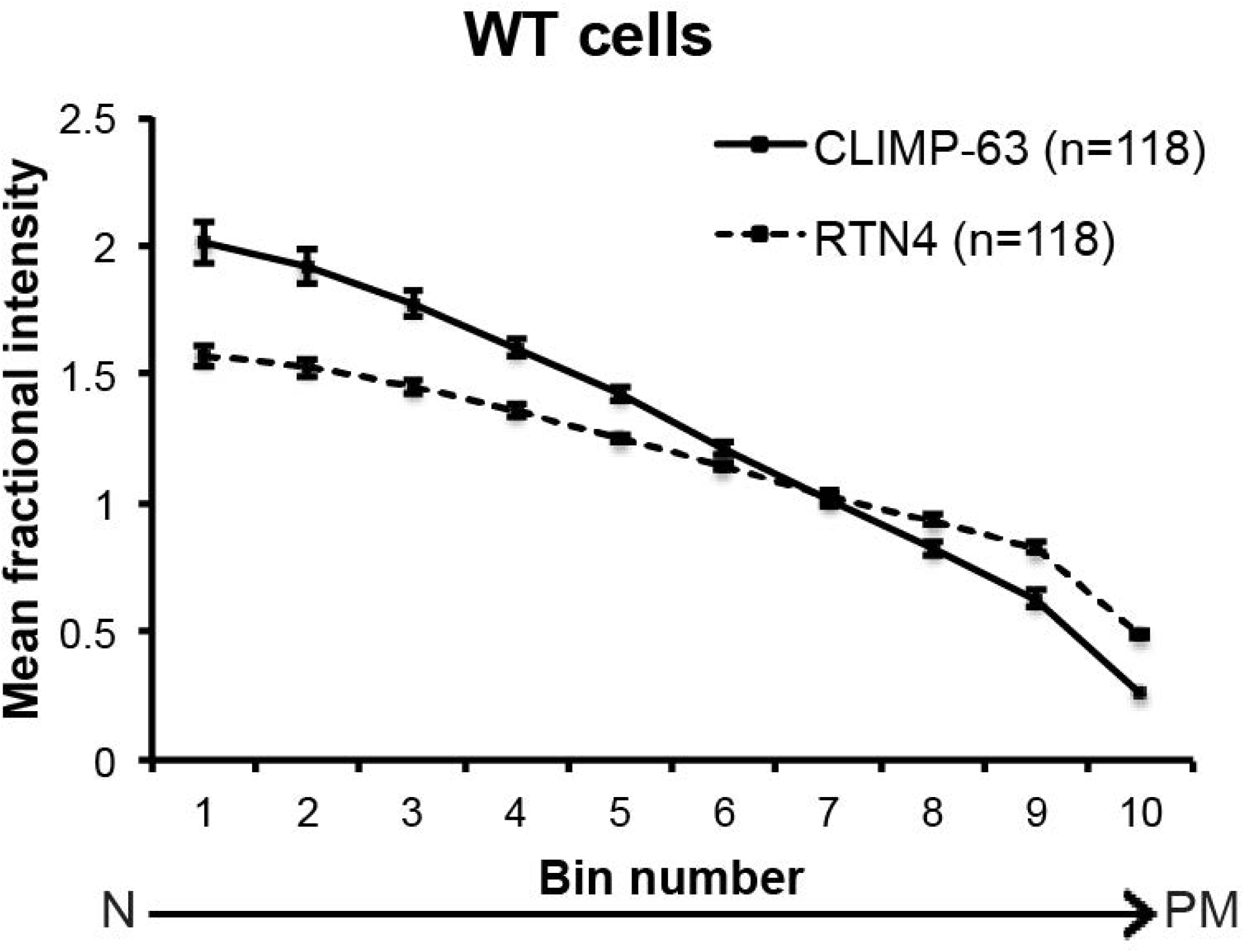
ER distribution in HeLa cells. Quantification of CLIMP-63 and RTN4 distribution in HeLa cells. Error bars represent the 95% CI. Representative immunofluorescence images are shown in Figure 3a. The results are representative of at least three independent experiments.

**Figure S4.**
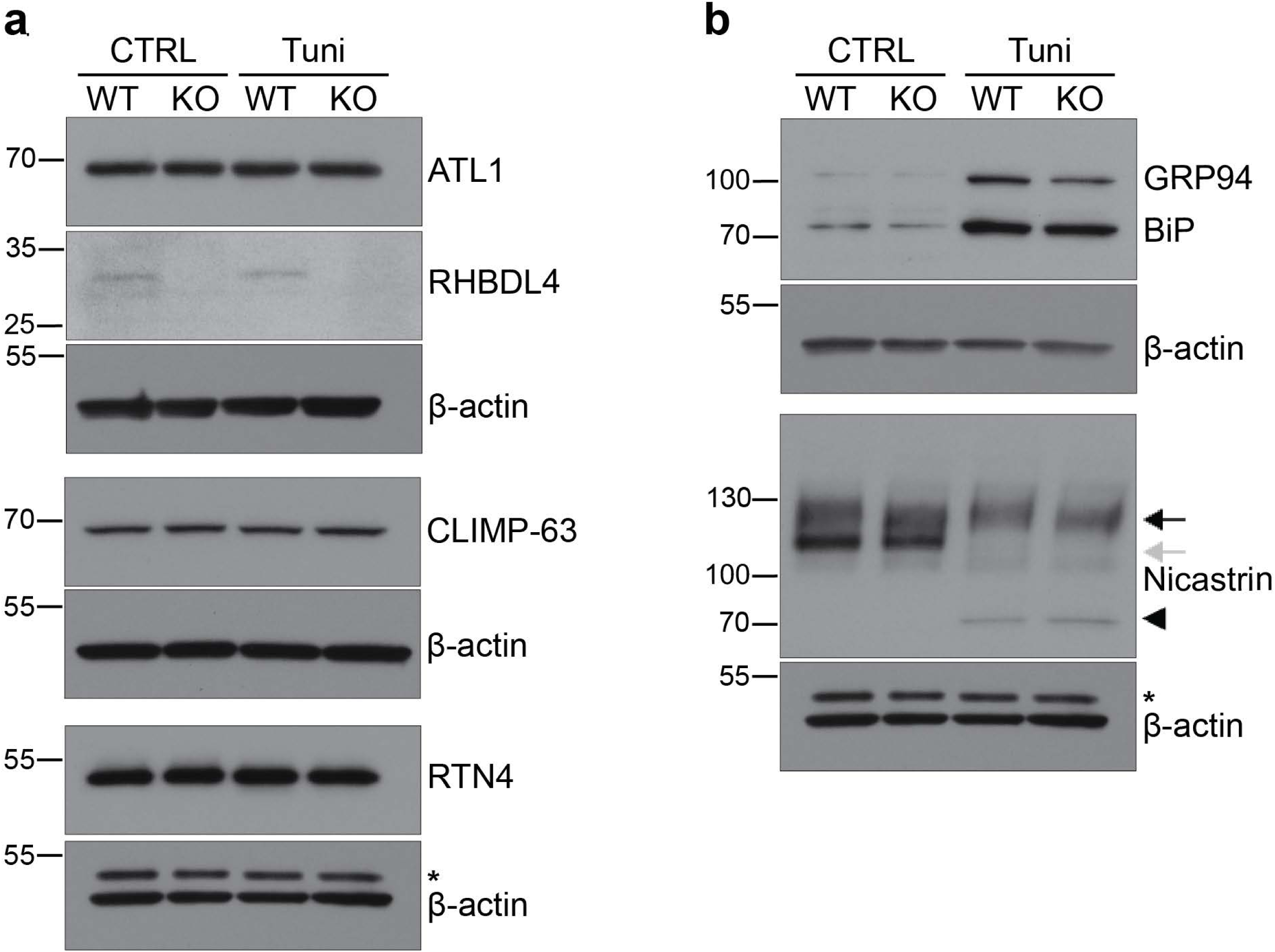
The levels of ER shaping proteins and the ER stress response are similar in WT and RHBDL4 KO cells. Western blot of total cell lysates from WT and RHBDL4 KO HeLa in control conditions or 24 hours after tunicamycin treatment. **a,** Western blots of RHBDL4 and ER shaping proteins ATL1, CLIMP-63 and RTN4. **b,** Western blots of UPR targets BiP and GRP94 (detected using an anti-KDEL antibody), and Nicastrin. The black arrow represents the mature post-Golgi protein, while the grey arrow represents the immature, ER-localised one. The arrowhead represents the non-glycosylated Nicastrin, visible in the tunicamycin-treated samples. β-actin was used as a loading control and it was developed on the stripped membranes, previously used for the proteins above each β-actin panel. The asterisk represents incompletely removed RTN4. The results are representative of three independent experiments.

**Figure S5.**
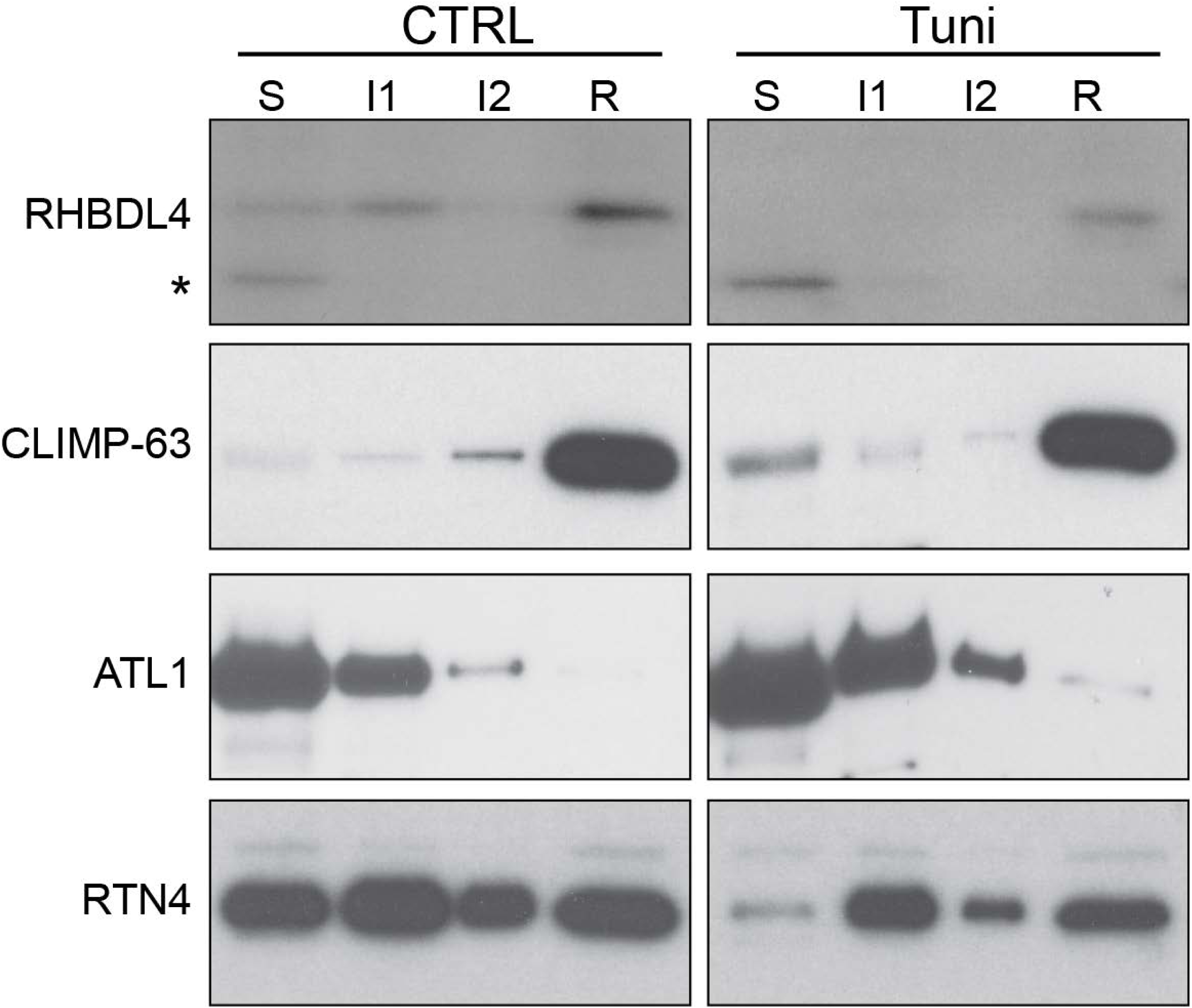
RHBDL4 resides within the ER sheets. Microsomes from MEFs, control and tunicamycin treated, were separated as in Figure 5a. Smooth ER (S), rough ER (R) and intermediate ER were analysed by western blot for the indicated ER proteins. The asterisk represents a non-specific band. The results are representative of three independent experiments.

**Figure S6.**
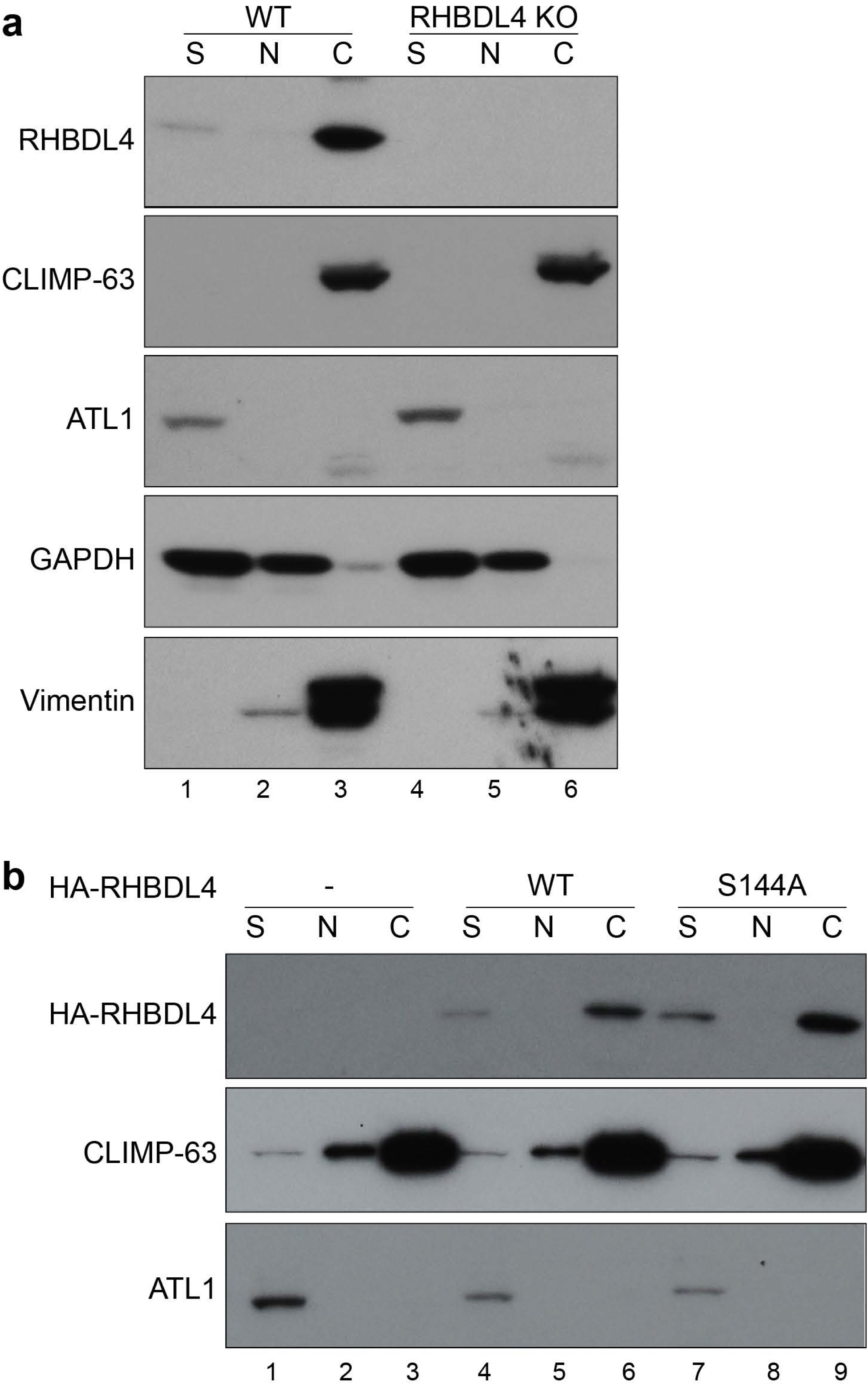
RHBDL4 associates with the cytoskeleton. **a,** Western blot of soluble (S), nuclear (N) and cytoskeletal (C) fractions isolated from WT and RHBDL4 KO MEFs and analysed for ER proteins – RHBDL4, CLIMP-63, ATL1 – as well as for GAPDH and vimentin. **b**, Anti-HA tag western blot of C, N and S fractions isolated from RHBDL4 KO MEFs transiently expressing HA-tagged RHBDL4 WT or RHBDL4 S144A. The results are representative of three independent experiments (**a**) or two independent experiments (**b**).

**Figure S7.**
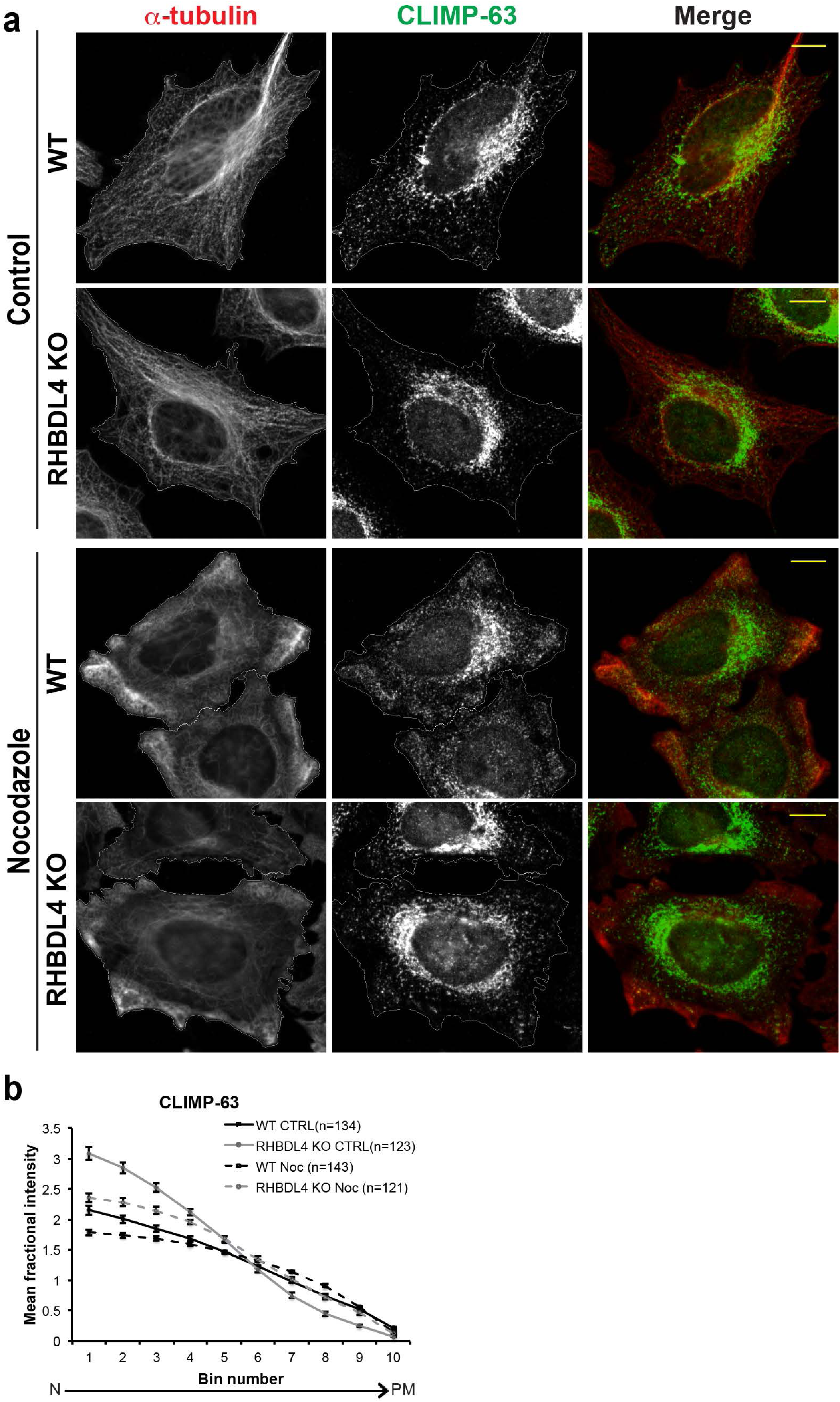
RHBDL4 knockout affects the ER-sheet re-distribution when microtubules are depolymerised. **a**, Immunofluorescence of WT and RHBDL4 KO HeLa cells showing the microtubule (α-tubulin – red) and ER-sheet (CLIMP-63 – green) distribution under control or nocodazole treatment. **b**, Quantification of CLIMP-63 distribution. Error bars represent the 95% CI. The results are representative of three independent experiments.

**Figure S8.**
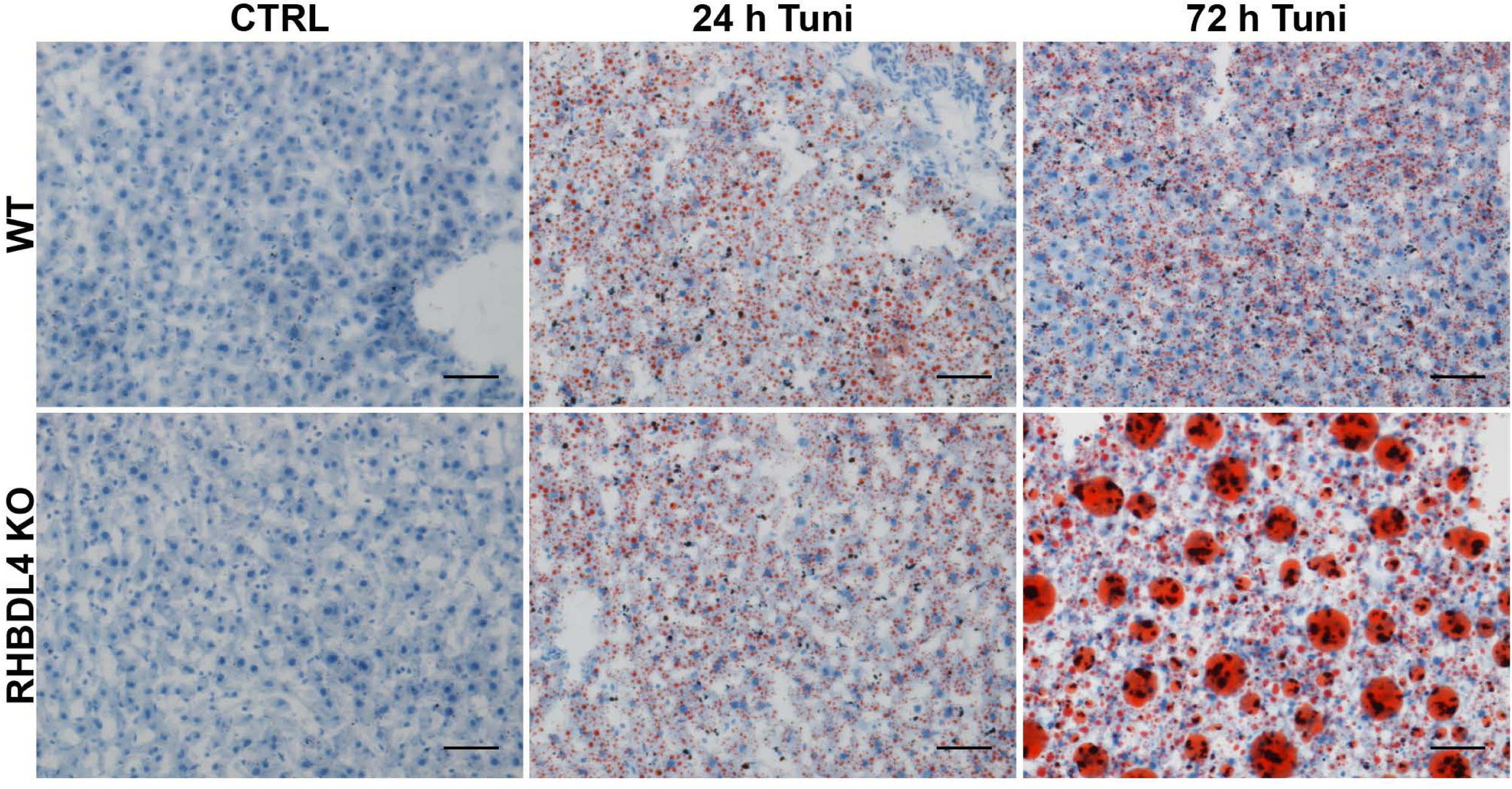
RHBDL4 protects against the ER stress in mice. Oil red O staining of liver tissue sections from WT and RHBDL4 KO mice, control, 24 or 72 hours after tunicamycin treatment. The images are representative of one experiment with n=3 mice per genotype for 24 hours treatment and n=2 mice per genotype for 72 hours treatment. n=4 mice per genotype for control.

## Notes

### Competing Interest Statement

The authors have declared no competing interest.

### Summary of Updates

Authorship updated

